# Tumor-intrinsic PRC2 inactivation drives a context-dependent immune-desert tumor microenvironment and confers resistance to immunotherapy

**DOI:** 10.1101/2022.05.27.493507

**Authors:** Juan Yan, Yuedan Chen, Amish J. Patel, Sarah Warda, Briana G. Nixon, Elissa W.P. Wong, Miguel A. Miranda-Román, Cindy J. Lee, Ning Yang, Yi Wang, Jessica Sher, Emily Giff, Fanying Tang, Ekta Khurana, Sam Singer, Yang Liu, Phillip M. Galbo, Jesper L. Maag, Richard P. Koche, Deyou Zheng, Liang Deng, Cristina R. Antonescu, Ming Li, Yu Chen, Ping Chi

**Affiliations:** Human Oncology and Pathogenesis Program, Memorial Sloan Kettering Cancer Center, New York, NY, USA; Weill Cornell Graduate School of Medical Sciences, Cornell University, New York, NY, USA; Immunology Program, Sloan Kettering Institute, Memorial Sloan Kettering Cancer Center, New York, NY, USA; Louis V. Gerstner, Jr., Graduate School of Biomedical Sciences, Memorial Sloan Kettering Cancer Center, New York, NY, USA; Dermatology Service, Department of Medicine, Memorial Sloan Kettering Cancer Center, New York, NY, USA; Department of Physiology and Biophysics, Weill Cornell Medicine, New York, NY, USA; Meyer Cancer Center, Weill Cornell Medicine, New York, NY, USA; Institute for Computational Biomedicine, Weill Cornell Medicine, New York, NY, USA; Department of Surgery, Memorial Sloan Kettering Cancer Center, New York, NY, USA; Department of Genetics, Albert Einstein College of Medicine, Bronx, NY, USA; Center for Epigenetics Research, Memorial Sloan Kettering Cancer Center, New York, NY, USA; Department of Neurology, Albert Einstein College of Medicine, Bronx, NY, USA; Department of Neuroscience, Albert Einstein College of Medicine, Bronx, NY, USA; Weill Cornell Medical College, New York, NY, USA; Department of Pathology, Memorial Sloan Kettering Cancer Center, New York, NY, USA; Department of Medicine, Memorial Sloan Kettering Cancer Center, New York, NY, USA

## Abstract

Immune checkpoint blockade (ICB) has demonstrated clinical success in “inflamed” tumors with significant T-cell infiltrates, but tumors with an immune-desert tumor microenvironment (TME) fail to benefit. The tumor cell-intrinsic molecular mechanisms of the immune-desert phenotype remain poorly understood. Here, we demonstrate that inactivation of the Polycomb-repressive complex 2 (PRC2) core components, EED or SUZ12, a prevalent genetic event in malignant peripheral nerve sheath tumor (MPNST) and sporadically in other cancer types, drives a context-dependent immune-desert TME. PRC2 inactivation reprograms the chromatin landscape that leads to a cell-autonomous shift from primed baseline signaling-dependent cellular responses (e.g., interferon γ) to PRC2-regulated development and cellular differentiation transcriptional programs. Further, PRC2 inactivation reprograms the TME, leads to diminished tumor immune infiltrates and immune evasion through reduced chemokine production and impaired antigen presentation and T-cell priming, and confers ICB primary resistance through blunted T-cell recruitment *in vivo*. We demonstrate that strategies that enhancing innate immunity via intratumoral delivery of inactivated modified vaccinia virus Ankara (MVA) leads to increased tumor immune infiltrates and sensitizes PRC2-loss tumors to ICB. Our results provide novel molecular mechanisms of context-dependent dysfunctional epigenetic reprogramming that underline the immune-desert phenotype in MPNST and other cancers with PRC2 inactivation. Importantly, our findings highlight genetic-inactivation of PRC2 as a novel context-dependent ICB therapeutic resistance biomarker in cancer, and caution that therapeutic strategies that non-selectively target PRC2 in the host may lead to undesirable context-dependent immune evasion and ICB resistance in tumors. Our studies also point to intratumoral delivery of immunogenic therapeutic viruses as an initial strategy to modulate the immune-desert TME and capitalize on the clinical benefit of ICB.

## Introduction

The PRC2 complex consisting of core components of EZH1/2, EED, and SUZ12, establishes and maintains H3K27me2/3 in the genome and regulates chromatin structure, transcription, cellular stemness, and differentiation [1]. PRC2 is a context-dependent tumor suppressor whose core components are frequently inactivated genetically or epigenetically in various cancer types, including MPNST [2, 3], melanoma [2], myeloid disorders [4, 5], T-cell acute lymphocytic leukemia (ALL) [6], and early T-cell precursor ALL [7], pediatric gliomas [8–11], invasive breast cancer [12], and others. Among all cancer types, high-grade MPNST, a group of aggressive soft tissue sarcomas with no effective therapies, has the highest prevalence of complete loss of PRC2 function through biallelic inactivation of the PRC2 core components, EED or SUZ12 [2, 3, 13–15].

There is increasing evidence demonstrating the critical context-dependent role of PRC2 in regulating immune cell identity and function, including cytotoxic CD8^+^ T cell repression [16–21], CD4^+^ T helper cell repression [22–24], and Treg activation [25–27] through regulating cell-lineage specific gene expression. Therefore, selectively targeting PRC2 in immune cells may modulate the TME and the tumor responses to immunotherapy [28]. Beyond immune cells, PRC2 in cancer cells has recently been shown to maintain bivalency at MHC-I antigen processing genes silencing MHC-I expression in selective MHC-I low cancer (e.g., SCLC, neuroblastoma) and targeting PRC2 can potentially enhance anti-tumor immunity through increasing MHC class I-antigen presentation in this setting [29]. However, it remains unclear how tumor-intrinsic PRC2 inactivation affects the tumor immune microenvironment, and whether PRC2 has similar regulation of MHC-I in MHC-I high cancer.

By utilizing both human MPNST tissues and engineered PRC2-loss mouse models in this study, we demonstrated that tumor-intrinsic PRC2-loss drives immune evasion through epigenetic reprogramming, and consequently, deficiency in antigen presentation, chemokine production, and IFNγ signaling, and primary resistance to ICB. To overcome the “cold” TME in PRC2-loss cancer, we used intratumoral delivery of immunogenic MVA and demonstrated that the inactivated MVA can enhance tumor immunity, alter the immune-desert TME, and sensitizes the PRC2-loss tumors to ICB therapy. These studies indicate that genetic inactivation of PRC2 can be used as a novel biomarker for resistance ICB therapy in selective cancers and that therapeutic viruses that can enhance tumor immunity maybe a potential first step to overcome the “cold” TME in PRC2-loss tumors.

## Results

### Tumor-intrinsic PRC2 loss is associated with immune-desert tumor microenvironment in MPNST

To characterize the role of PRC2 inactivation in cancer pathogenesis, we analyzed the transcriptomes of 41 histologically confirmed high-grade human MPNST tumor samples, consisting of both PRC2-wild-type (wt) and PRC2-loss samples. PRC2 loss in MPNSTs was confirmed by loss of H3K27me3 immunostaining and/or genetic inactivation of EED or SUZ12 by MSK-IMPACT [30] (**Table S1**). Principal component analysis (PCA), which detects sources of variation, showed that the MPNST samples were readily separated by PRC2 status in the first principal component (PC1) (**Fig. S1A**). We generated a gene set composed of genes that were differentially expressed between PRC2-loss and PRC2-wt samples. Hierarchical clustering of these genes robustly separated the PRC2-loss and PRC2-wt MPNSTs, with the majority of the most differentially expressed genes upregulated in PRC2-loss compared to PRC2-wt MPNSTs, consistent with the role of PRC2 in transcriptional repression (**Fig. 1A**). Consistently, gene set enrichment analysis (GSEA) of these genes showed that the most enriched pathways and gene sets in PRC2-loss MPNSTs included the PRC2 modules and H3K27me3 target genes, organ development and morphogenesis, neuron cell fate specification and WNT signaling gene sets [3, 31–33] (**Fig. 1B, S1B, Table S2**). A distinct smaller subset of genes was consistently downregulated in PRC2-loss compared to PRC2-wt MPNSTs (**Fig. 1A**). Remarkably, nearly all these genes were associated with immune function, including both innate and adaptive immune response pathways, T- and B-cell receptor signaling pathways, antigen binding and presentation (**Fig. 1B, Table S2**).

**Figure 1.**
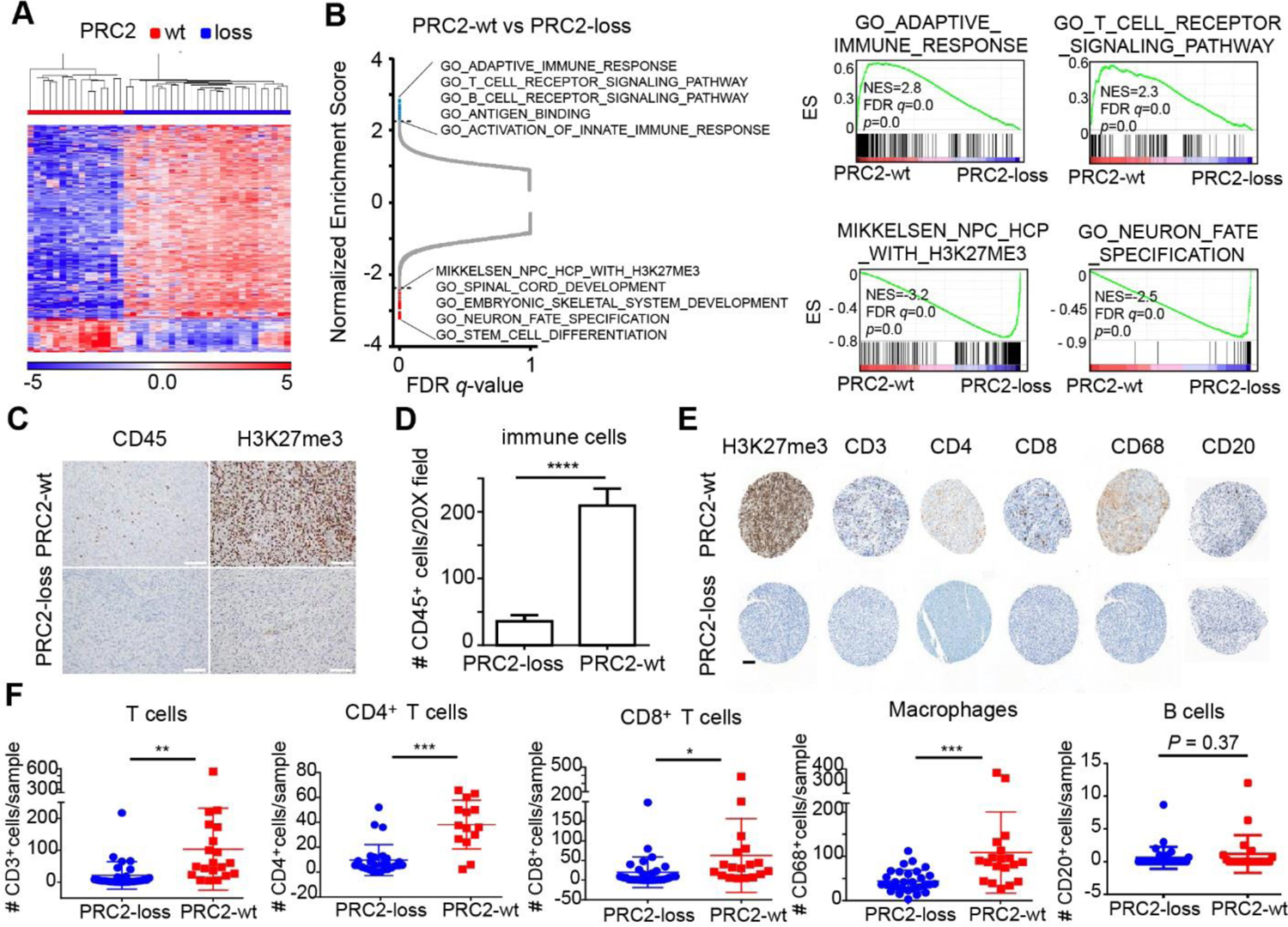
PRC2 loss is associated with deficiency of broad subclasses of tumor immune infiltrates in MPNST. **A**, Hierarchical clustering of the most differentially regulated genes (FDR *q* ˂ 0.05, and fold-change ≥ 8) between PRC2-loss and PRC2-wt MPNST tumors by RNA-seq analysis. **B**, GSEA of PRC2-wt vs. PRC2-loss MPNST transcriptomes. Left: GSEA normalized enrichment score (NES) vs. FDR *q*-value. Blank dash line: top 100 gene sets; Red dots: PRC2 and development related gene sets; Blue dots: immune related gene sets. Right: representative GSEA profiles of negatively enriched and positively enriched gene sets with PRC2-loss. **C**, Representative IHC of H3K27me3 and CD45 in PRC2-wt and PRC2-loss MPNST tumors. Scale bar: 100 μm **D**, Quantification of CD45^+^ cells in PRC2-wt (n=39) and PRC2-loss (n=69) MPNST tumors. \*\*\*\**P*<0.0001 by unpaired two-tailed t test. Error bars: mean±SEM. **E**, Representative IHC of H3K27me3, CD3, CD4, CD8, CD68 and CD20 in PRC2-wt (n=21) and PRC2-loss (n=30) tumors from MPNST tissue microarray. Scale bar: 100 μm. **F**, Quantification of CD3^+^ T cells, CD4^+^ T cells, CD8^+^ T cells, CD68^+^ macrophages/monocytes and CD20^+^ B cells in PRC2-wt and PRC2-loss MPNST TMA tumors. \**P*<0.05, \*\**P*<0.01, \*\*\**P*<0.001 by unpaired two-tailed t test. Error bars: mean±SD.

We next performed immunostaining of several established markers of distinct immune subclasses by immunohistochemistry (IHC) in human MPNST tissue microarrays (TMA), including CD45 (pan-leukocyte), CD3 (T cells, CD4^+^ and CD8^+^ subsets), CD68 (macrophages/monocytes) and CD20 (B cells). Quantification of each immune subset revealed that compared to PRC2-wt, the PRC2-loss MPNSTs were associated with significant reductions in CD45^+^ leukocytes (**Fig. 1C-1D**), CD3^+^ T cells, CD4^+^ and CD8^+^ T cells, and CD68^+^ macrophages/monocytes (**Fig. 1E-1F**). Although not significantly different, CD20^+^ B cells were only marginally present in both PRC2-wt and PRC2-loss MPNSTs (**Fig. 1E-1F**). These data corroborate the transcriptome results of diminished tumor immune infiltrates (**Fig. S1C**) and suggest that tumor cell-intrinsic PRC2 inactivation may exclude immune infiltrates and drive an immune-desert TME.

### Antigen presentation and IFNγ signaling are suppressed in PRC2-loss MPNST

To understand how tumor-intrinsic PRC2 loss leads to an immune-desert TME, we further characterized the transcriptomes of PRC2-loss vs. PRC2-wt MPNST tumors and focused on the initiating steps of anti-tumor immune response [34]. Among the genes downregulated by PRC2 loss, antigen processing and presentation was one of the most negatively enriched gene sets by GSEA (**Fig. 2A, Table S2**). Interferon gamma (IFNγ) is an established key regulator of antigen presentation and chemokines for immune cell recruitment [35, 36]. Consistently, IFNγ signaling was significantly impaired in PRC2-loss tumors, including cellular response to IFNγ, interferon responsive genes and regulation of IFNγ production (**Fig. 2A**). The expression of the tumor cell-autonomous genes, *IFNGR1* and downstream signaling axis and effectors, including *JAK1*, *JAK2*, *IRF1*, were all significantly lower in PRC2-loss compared to PRC2-wt tumors (*P* < 0.05) (**Fig. 2B**). In addition, IFNγ, usually produced by immune cells, was very low in most PRC2-loss tumors (**Fig. 2B**), consistent with PRC2-loss associated immune-desert phenotype. Overall, the expression of the IFNG-IFNGR1-JAK signaling axis genes was significantly reduced in PRC2-loss tumors compared to PRC2-wt. Consequently, genes relevant to antigen processing and presentation were also decreased accompanying PRC2 loss, including MHC class I (e.g., *HLA-A* and *B2M*), MHC class II (e.g., *CD74* and *HLA-DMA*) and antigen processing genes (e.g., *TAP1*) (**Fig. 2B**), suggesting decreased tumor immunogenicity in PRC2-loss tumors compared to PRC2-wt. We further validated the decreased protein expression of MHC class I, B2M and MHC class II by IHC in PRC2-loss tumors using MPNST TMA (**Fig. 2C-2D**). Furthermore, the key chemokines responsible for immune cell recruitment were significantly lower in PRC2-loss compared to PRC2-wt tumors, including *CXCL9/10* (*P* < 0.01) and *CCL2/3/4/5* (*P* < 0.05) (**Fig. 2B**). These data further posit that the tumor cell-intrinsic PRC2 loss-associated immune-desert phenotype is driven by impaired antigen presentation and diminished IFNγ signaling in tumors.

**Figure 2.**
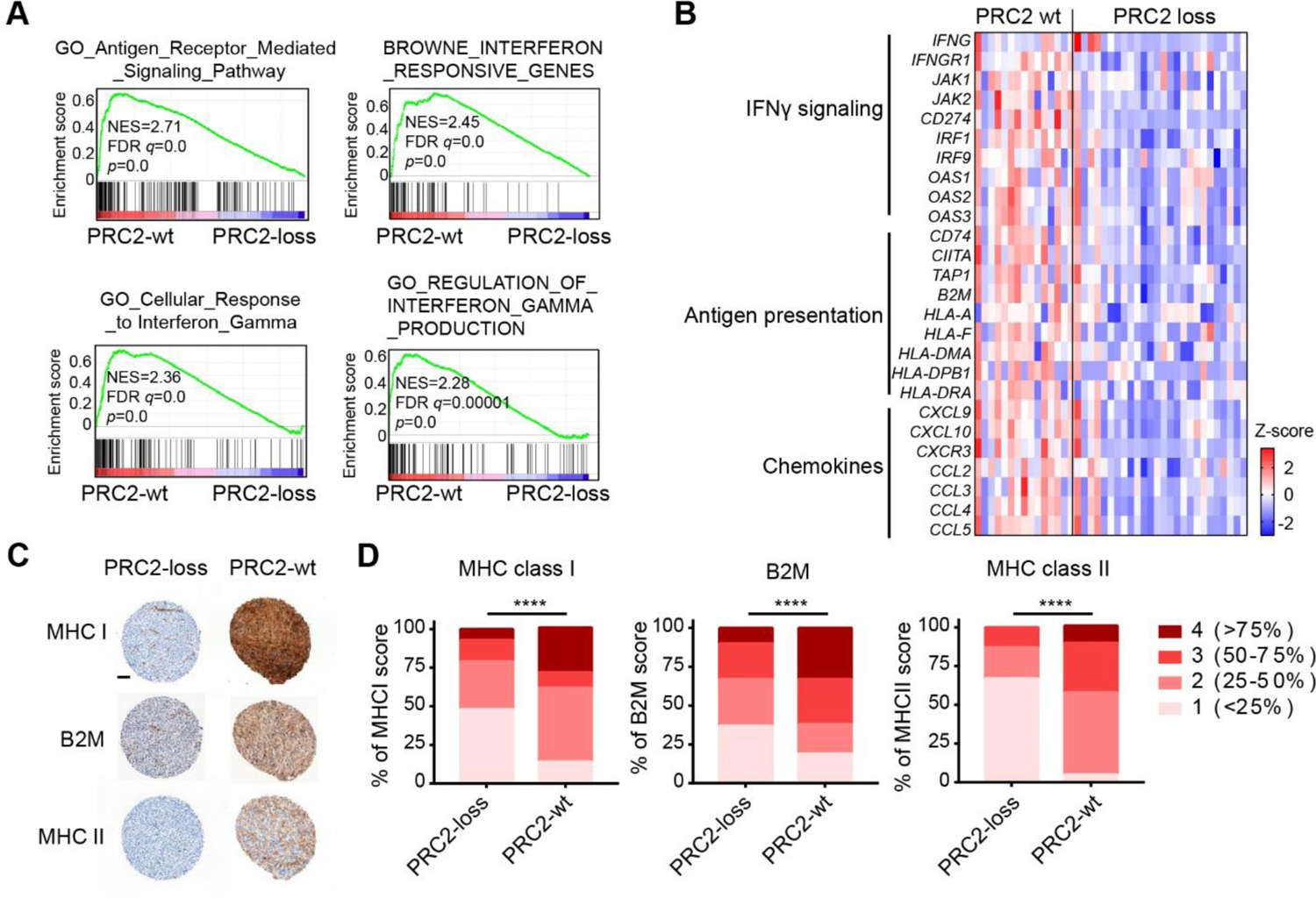
Antigen presentation and IFN γ pathway are diminished in PRC2-loss MPNST. **A**, GSEA of human MPNST transcriptomes points to antigen presentation and IFNγ pathway defects in PRC2-loss compared to PRC2-wt tumors. **B**, Heatmap of IFNγ signaling and downstream genes in human MPNSTs. **C**, Representative IHC of MHC I, MHC II and B2M in PRC2-wt and PRC2-loss human MPNST tumor samples. Scale bar: 100 μm. **D**, Quantification of MHC I, MHC II, and B2M in PRC2-loss and PRC2-wt human MPNSTs. Chi-square test for contingency, \*\*\*\**P*<0.0001

### PRC2 loss reprograms the chromatin landscape and suppresses subset of IFN-targeted genes

To examine the role of PRC2 loss in MPNST while minimizing cell line-specific confounding factors, we generated and validated PRC2-isogenic human MPNST cells using CRISPR/Cas9-mediated knockout of the PRC2 core component, SUZ12, in a PRC2-wt, *NF1^-/-^; CDKN2A^-/-^* M3 cell line derived from a human NF1-associated MPNST (**Fig. 3A**). SUZ12 loss led to a global reduction of the H3K27me3, H3K27me2 and H3K27me1 marks, and a reciprocal global increase of the H3K27ac mark in PRC2-isogenic human MPNST cells (sg*Con* vs. sg*SUZ12*) (**Fig. 3A**). The PRC2-isogenic M3 cells, when orthotopically transplanted into the sciatic nerve pockets of immunodeficient NSG mice, gave rise to high-grade MPNSTs with histological (e.g., monotonous spindle cell morphology, herringbone pattern and fascicular growth) and immunostaining (e.g., H3K27me3 and Ki67 staining) features resembling those of high-grade human MPNST [13] (**Fig. 3A**). Nucleocytoplasmic and chromatin fractionation demonstrated that the global decrease of H3K27me3 and increase of H3K27ac occurred on chromatin in SUZ12-loss cells (**Fig. 3B**). These characterizations combined with the loss of H3K27me3 immunostaining in PRC2-loss M3 cell line-derived MPNST tumors validate the model system for mechanistic studies.

**Figure 3.**
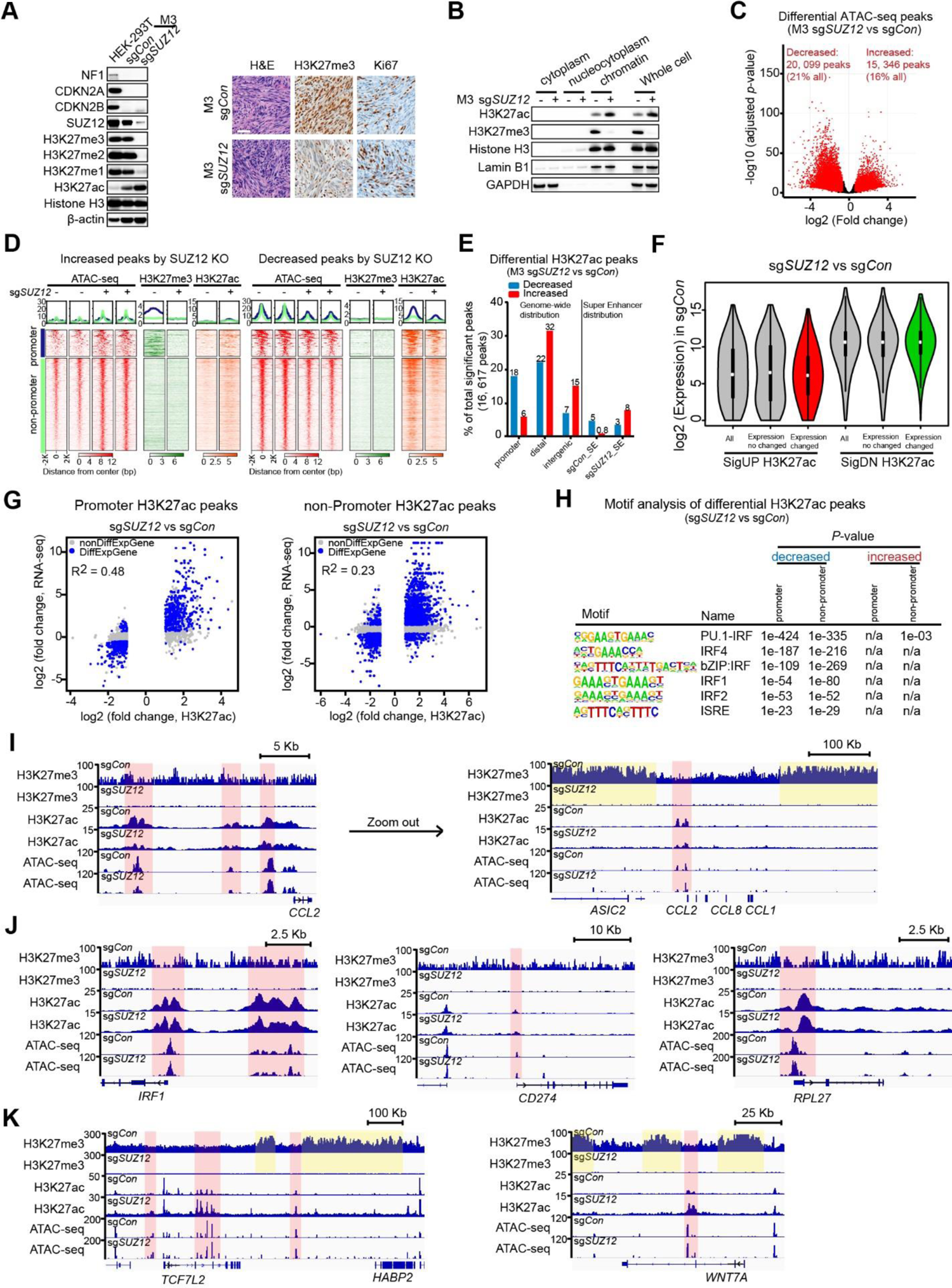
PRC2 loss reprograms the chromatin landscape and partially suppresses IFNγ signaling. **A**, Representative immunoblots and IHC of indicated proteins and histone modifications in PRC2-loss (sg*SUZ12*) and PRC2-wt (sg*Con*) isogenic human MPNST cells (left) and orthotopically (sciatic nerve pocket) transplanted MPNST tumors (right). Scale bar: 50 μm. **B**, Representative immunoblots of indicated histone modifications and control proteins from the cytoplasm, nucleocytoplasm and chromatin fractions of PRC2-isogenic M3 cells. **C**, A volcano plot of chromatin accessibility changes by ATAC-seq comparing PRC2-loss (sg*SUZ12*) with PRC2-wt (sg*Con*) isogenic M3 cells. Red dots represent significantly changed ATAC peak (FDR *q* ˂ 0.1, fold-change ≥ 1.5). **D**, Density plot of various histone modifications by ChIP-seq, centered on significantly increased (left) and decreased ATAC-peaks (right) in PRC2-loss (sg*SUZ12*) compared to PRC2-wt (sg*Con*) isogenic M3 cells. Promoter: transcriptional start site (TSS) ± 2kb; Non-promoters: rest of the genome other than promoter, including distal regulatory enhancer and intergenic regions. **E**, Distribution of PRC2 loss-associated significantly decreased (blue) and increased (red) H3K27ac peaks across different genomic regions in PRC2-isogenic M3 cells (FDR *q* ˂ 0.05, fold-change ≥ 2); promoter (TSS ± 2 kb), distal regulatory (−50 kb from TSS to transcriptional end site [TES] + 5 kb) and intergenic (non-promoter, non-distal regulatory) regions, as well as at super enhancers (SE). **F**, Violin plots of mRNA baseline expression of genes associated with PRC2-loss induced significantly increased (SigUP) and decreased (SigDN) H3K27ac enrichment at their respective loci in in PRC2-wt (sg*Con*) M3 cells. **G**, The correlation of significant transcriptome and H3K27ac changes at promoter (TSS ± 2kb) and non-promoter regions (FDR *q* < 0.05, fold-change ≥ 2). Blue and grey dots represent peaks mapped to genes with (DiffExpGene) and without (nonDiffExpGene) significant transcriptome changes, respectively. **H**, HOMER motif analysis of PRC2 loss-associated significantly decreased and increased H3K27ac peaks in PRC2-isogenic M3 cells, focusing on comparison of immune response related motifs. **I-K**, ChIP-seq and ATAC-seq profiles at the loci of selective IFNγ-responsive genes, e.g., *CCL2* (I), *IRF1* (J), *CD274* (J), control gene *RPL27* (J), *TCF7L2* (K) and *WNT7A* (K) in PRC2-isogenic M3 cells. Pink highlights regions with H3K27ac enrichment peaks, while yellow highlights regions with high H3K27me3 enrichment in sg*Con*.

We speculated that PRC2 loss may alter the chromatin context and directly affect the transcriptional regulation of genes related to immune signaling and responses. We first examined the impact of PRC2 loss on genome-wide distribution of chromatin accessibility by ATAC-seq and PRC2 relevant chromatin marks (H3K27me3 and H3K27ac) by ChIP-seq, in PRC2-isogenic M3 cells. Globally, compared to PRC2-wt, PRC2 loss led to not only significant increase, but also significant decrease of chromatin accessibility at 15,346 (16% of all ATAC peaks) and 20,099 genomic loci (21% of all ATAC peaks), respectively (**Fig. 3C, S2A**). The significantly changed chromatin accessibility regions (increased or decreased) were relatively similarly distributed at promoter (7.7%, 8.3%) and non-promoter regions, including distal regulatory (59.9%, 68.8%) and intergenic regions (32.4%, 22.9%) (**Fig. S2B)**. The significantly increased ATAC peaks associated with PRC2 loss overlapped more with H3K27me3 enriched loci (10.5%) than the decreased ATAC peaks (2.4%) (**Fig. S2C**), consistent with the PRC2 function on chromatin compaction and transcription. Importantly, the increased ATAC peaks localized to both H3K27me3 enriched promoter and non-promoter distal regulatory and intergenic regions devoid of H3K27me3 in PRC2-wt controls and were associated with a modest reciprocal increase of H3K27ac enrichment at both promoters and non-promoters with PRC2 loss (**Fig. 3D, S2D).** In contrast, the genomic regions with significant decrease in chromatin accessibility by PRC2 loss had minimal enrichment of H3K27me3 in PRC2-wt controls and were associated with decreased enrichment of H3K27ac at promoters and distal regulatory regions (**Fig. 3D, S2E**).

H3K27ac is an established chromatin mark preferentially enriched at active promoters and distal regulatory enhancers [13, 37, 38]. Despite the increase of total amount of chromatin bound H3K27ac with PRC2 loss, differential analysis of H3K27ac enrichment peaks showed relatively balanced gains and losses in the genome (**Fig. 3E**). While the significantly gained H3K27ac peaks mainly localized to distantly regulatory and intergenic regions and overlapped with H3K27me3 peaks, the significantly lost H3K27ac peaks mainly localized to promoters and existing super enhancers (SE) in PRC2-wt control with minimal overlap with H3K27me3 (**Fig. 3E, S2F**). Notably, the significantly decreased H3K27ac peaks localized to genes with higher baseline expression levels compared to those of increased H3K27ac peaks (**Fig. 3F**). Moreover, H3K27ac changes were well correlated with transcriptome changes at both promoter and non-promoter regions (**Fig. 3G**). Motif analysis demonstrated significant enrichment of transcription factor binding motifs associated with immune signaling pathways and responses, e.g., interferon signaling associated IRF family and ISRE motifs, only in the decreased but not increased H3K27ac peaks associated with PRC2 loss (**Fig. 3H, Table S3**). A subset of IFNγ-responsive gene loci was directly affected by PRC2 loss. For example, the H3K27ac enrichment at the super enhancer locus of the monocyte chemotactic protein, *CCL2*, was significantly diminished by PRC2 loss (sg*SUZ12*) compared to controls (**Fig. 3I**), whereas the H3K27ac enrichment at other IFNγ-target gene loci (e.g., *IRF1*, *CD274*) was unchanged (**Fig. 3J**). We observed that many lost and gained super enhancer regions by PRC2 loss were preferentially flanked by broad enrichment of H3K27me3 in the genome of PRC2-wt (**Fig. 3I, 3K**). These data suggest that PRC2 loss affects a broad region of H3K27ac enrichments and reprograms the genome-wide chromatin context of both the promoter and enhancer landscapes, which in turn alters signaling-dependent transcriptional response.

### Decreased chromatin accessibility for IFNγ-responsive loci in PRC2-loss MPNST cells

One of the fundamental functions of the chromatin context is to prime the cells for signaling dependent transcriptional response. Reasoning that altered local chromatin context mediated by PRC2 loss may change the transcriptional output, we next examined the transcriptome changes of IFNγ-responsive genes in response to IFNγ stimulation in PRC2-isogenic MPNST cells. Stimulation with 10 ng/ml exogenous IFNγ for 24 hours had no significant impact on the levels of H3K27me3 modification and *SUZ12*/*EED* mRNA expression in PRC2-isogenic M3 cells (**Fig. S3A-S3B**). PCA of ATAC-seq replicates under various conditions demonstrated robust clustering of replicates and separation of samples based on PRC2 status (PC1) and IFNγ stimulation (PC2) (**Fig. 4A**). *K*-means clusters 2 and clusters 6/7 were most representative of increased chromatin accessibility in response to IFNγ stimulation in the PRC2-loss (sg*SUZ12*) and PRC2-wt (sg*Con*) contexts, respectively (**Fig. 4B**). Consistently, *de novo* motif analysis of the *K*-means clusters showed that only clusters 2, 6 and 7 identified IFNγ stimulation related motifs (e.g. IRF8 and PU.1:IRF8) but with differential significance. The IFNγ-related motif IRF8 is the topmost enriched motif of cluster 7 with a *P* = 1e^-1039^, consist with a primed chromatin context in response to IFNγ stimulation through IRF activation in the PRC2-wt context. In contrast, the topmost enriched motif of cluster 2 is the GCN4/AP1 motif (*P* =1e^-1223^), whereas the classic IFNγ signaling relevant motif, IRF8/PU.1 is less significantly enriched (*P* =1e^-283^) (**Fig. 4B, Table S4**), suggesting a shift from IRF signaling to GCN4/AP1 signaling in a PRC2 loss-primed chromatin context. Moreover, GO analysis also revealed significant differences between the PRC2-loss representative cluster 2 and the PRC2-wt representative cluster 7 of chromatin accessibility changes induced by IFNγ. Cluster 7 was most enriched with immune response-related signaling pathways by GO analysis, including antigen presentation and interferon signaling (**Fig. 4C**), whereas cluster 2 was most enriched with development related pathways, e.g. cellular differentiation, organ development and PRC2 and H3K27me3 targets (**Fig. 4D**).

**Figure 4.**
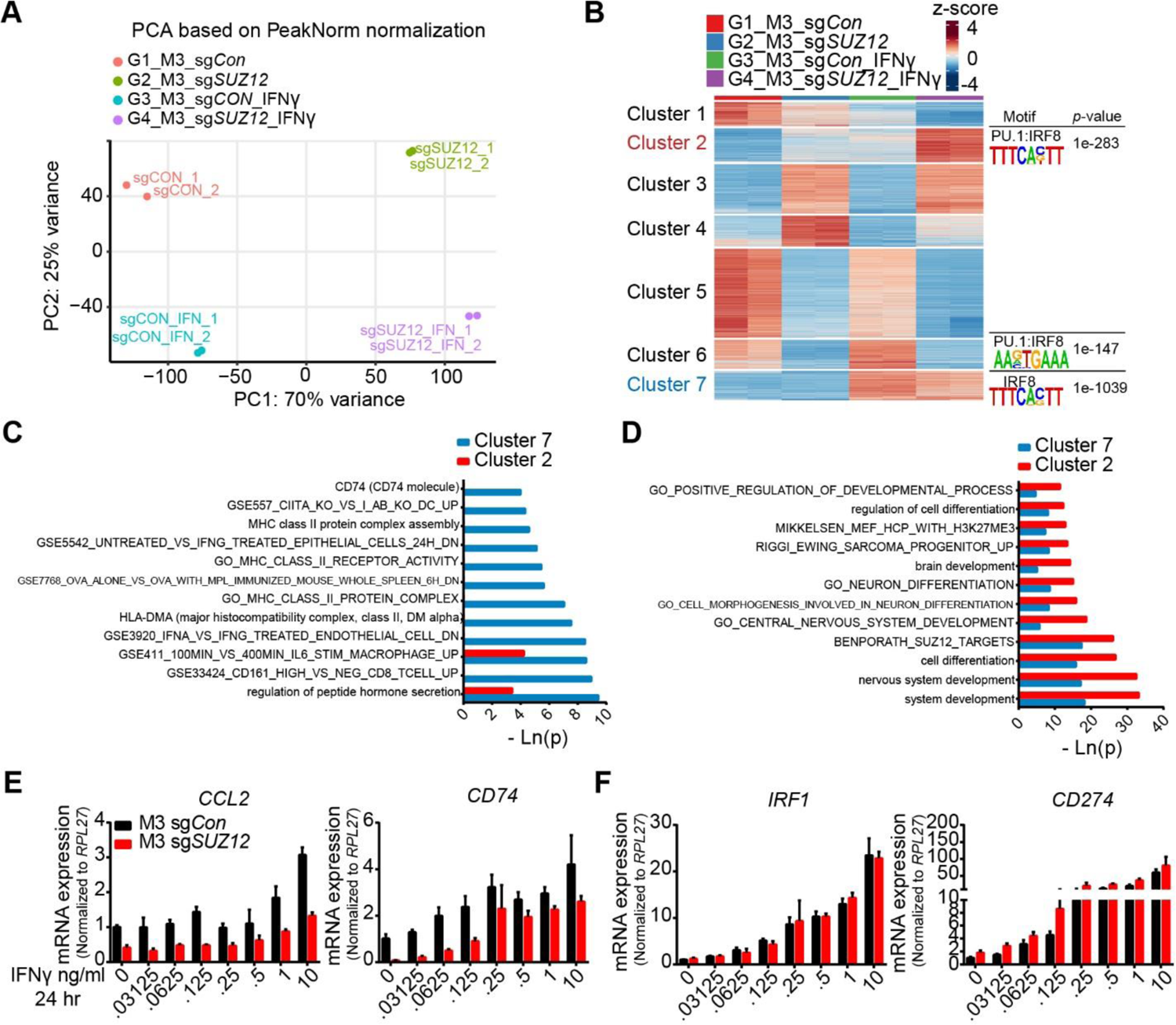
PRC2 loss dampens the IFNγ response in tumor cells through decreasing chromatin accessibility. **A**, Principal component analysis of chromatin accessibility by ATCA-seq in isogenic PRC2-wt (sg*Con*) and PRC2-loss (sg*SUZ12*) human MPNST cells with and without IFNγ stimulation. **B**, K-means clustering analysis of the chromatin accessibility changes in PRC2-isogenic M3 cells with or without IFNγ stimulation. Cells were treated with or without 10 ng/ml IFNγ for 24 hr followed by ATAT-seq. Homer de novo motif analysis enriches IRF only in cluster 2, 6 and 7. **C-D**, Comparison of Go analysis between cluster 7 and cluster 2 related to (B). **E-F**, IFNγ dose-dependent mRNA expression changes by qRT-PCR of IFNγ-responsive genes with or without PRC2 loss-associated chromatin accessibility changes, e.g. *CCL2* and *CD74* (E), *IRF1* and *CD274* (F), respectively. n=2, technical replicates. Error bars: mean±SEM.

Nevertheless, the transcriptome change tracked the changes in local chromatin context. In response to IFNγ stimulation, the transcription of IFNγ-response genes was unchanged when the local chromatin context was unperturbed, e.g., *IRF1* and *CD274* (**Fig. 4F**); and the transcriptional activation of IFNγ-response genes was significantly blunted, when the gene loci were associated with a decrease in chromatin accessibility and H3K27ac signal as a result of PRC2 loss, e.g., *CCL2*, *CD74*, *CIITA* and *HLA-DRA* (**Fig. 4E, S2E, S3C**). Further, the blunted transcriptional activation of these genes was most pronounced when IFNγ source was low (**Fig. 4E, S3C**). These observations suggest that PRC2 loss reprograms the steady-state primed chromatin context and leads to a blunted IFNγ response in tumor cells.

### Engineered PRC2 loss recapitulates the immune evasion through diminished antigen presentation and IFNγ signaling in both MPNST and breast cancer murine model

To evaluate the impact of tumor cell-intrinsic PRC2 inactivation on TME *in vivo*, we generated a histologically confirmed *Nf1^-/-^; Cdkn2a/b^-/-^* murine MPNST tumor-derived cell line (SKP605) from skin-derived precursors (SKPs) of C57BL/6J mice [39, 40] (**Fig. S4A-S4B**). Using CRISPR/Cas9-mediated knockout of PRC2 core component, *Eed*, we generated PRC2-isogenic murine MPNST cells (SKP605, sg*Con* vs. sg*Eed*) amenable for orthotopic and syngeneic transplant in immunocompetent C57BL/6J mice (**Fig. 5A-5B**). The PRC2-loss (sg*Eed*) orthotopically transplanted tumors exhibited accelerated growth in the sciatic nerve pockets of C57BL/6J mice compared to those of PRC2-wt (sg*Con*) tumors (**Fig. 5C-5D, S4C**). The expressions of immune cell recruitment chemokines e.g., *Ccl2* and *Cxcl10*, and lymphocyte activation cytokine, *Il2*, were significantly decreased in PRC2-loss tumors compared to PRC2-wt (**Fig. S4D**), indicating suppressed immune cell infiltration. Next, profiling of tumor immune infiltrates by FACS demonstrated a significant reduction of CD45^+^ leukocytes in PRC2-loss tumors compared to PRC2-wt (**Fig. 5E, S4E**). Further, the reduction of the tumor immune infiltrates in PRC2-loss tumors was across all major subclasses of immune cells, including MHCII^+^CD11c^+^ dendritic cells (DCs), TCRβ^+^ T cells, B220^+^ B cells, and to a lesser extent F4/80^hi^CD11b^+^ macrophages (**Fig. 5F, S4F, S5**), phenocopying the TME of human PRC2-loss MPNST (**Fig. 1C-1F**). Importantly, the IFNγ^+^ and TNFα^+^ CD4^+^ T cells were both significantly reduced in PRC2-loss tumors (**Fig. 5G, S4G**). These data suggest that PRC2 inactivation in MPNST leads to an immunosuppressive TME through both diminished recruitment of tumor immune infiltrates and functional T cells.

**Figure 5.**
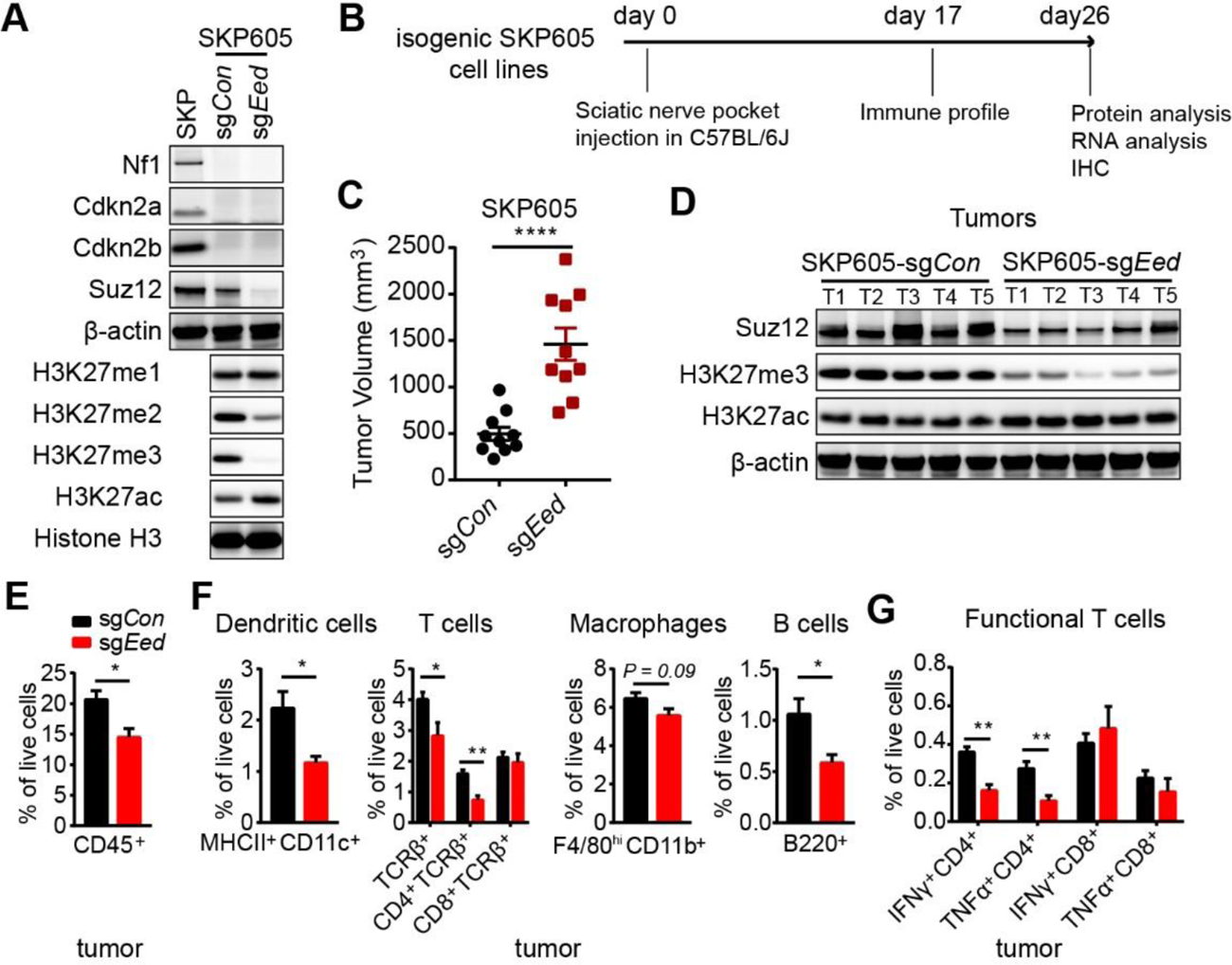
PRC2 loss suppresses tumor immune infiltration in a murine MPNST syngeneic transplant model. **A**, Representative immunoblots of indicated proteins and histone modifications in PRC2-loss (sg*Eed*) and PRC2-wt (sg*Con*) isogenic murine SKP605 MPNST cells. **B**, A schematic of the experimental and analysis plan of an orthotopic and syngeneic transplantable murine SKP605 MPNST model. **C**, Tumor volumes of PRC2-isogenic (sg*Eed* vs. sg*Con*) murine SKP605 tumors on day 26 post sciatic nerve pocket implantation. n=10 tumors bilaterally grafted in C57BL/6J mice for each PRC2-wt (sg*Con*) and PRC2-loss (sg*Eed*) cohort. **D**, Representative immunoblots of indicated proteins and histone modifications of selective PRC2-wt and PRC2-loss murine MPNST tumors in (C). **E-F**, Percentage of CD45^+^ (E) and subpopulations (F), including dendritic cells (B220^-^F4/80^lo^MHCII^+^CD11c^+^), T cells (TCRβ^+^), macrophages (B220^-^ F4/80^hi^CD11b^+^) and B cells (B220^+^), of total live cells in PRC2-isogenic murine SKP605 tumors in (C), n=5 tumors for each PRC2-wt (sg*Con*) and PRC2-loss (sg*Eed*) cohort. **G**, The percentage of IFNγ^+^CD4^+^ or IFNγ^+^CD8^+^ T cells of total live cells in the PRC2-isogenic SKP605 tumors. n=5 tumors for each group. C, and E-G: \**P*<0.05, \*\**P*<0.01, **** *P*<0.0001 by unpaired two-tailed t test. All error bars: mean±SEM.

To evaluate whether PRC2 loss exerts similar effect on tumor immune microenvironment in other cancer types, we used CRISPR/Cas9-mediated knockout of PRC2 core components, *Eed* or *Suz12*, and generated PRC2-isogenic murine mammary tumor model (AT3, sg*Con* vs. sg*Eed* or sg*Suz12*) amenable for syngeneic transplant in C57BL/6J mice. Although PRC2 loss did not affect the tumor cell growth *in vitro* (**Fig. S6A**), it accelerated orthotopically and syngeneically grafted tumor growth *in vivo* (**Fig. 6A-6B, S6B).** PRC2 downstream target H3K27me3 loss was maintained in grafted PRC2-loss tumors (**Fig. 6C, S6C**). Transcriptome analysis of the explanted PRC2-wt (sg*Con*) and PRC2-loss (sg*Eed*) AT3 tumors by RNA-seq demonstrated that PRC2 loss led to the upregulation of various developmental pathways, including Wnt/β-catenin signaling and PRC2/ H3K27me3 targets, and the downregulation of both innate and adaptive immune response pathways, including antigen processing and presentation and IFNγ response pathways (**Fig. 6D, Table S5**). Consistently, there was significant reduction of the expression of *Ifng* and Ifnγ signaling-related genes (e.g., *Ifngr1*, *Cd274*, *Irf1*, *Irf9*), antigen presentation-related genes (e.g., *Tap1*, *Tap2*), MHC class I (e.g., *H2-k1*, *H2-q4*, *H2-23*), and MHC class II (e.g., *H2-t10* and *H2-aa*) genes in PRC2-loss tumors compared to PRC2-wt (**Fig. 6E**), indicating impaired tumor immunogenicity by tumor cell-intrinsic PRC2 inactivation. Moreover, there were less immune recruitment chemokines in PRC2-loss tumors (e.g., *Cxcl9*, *Cxcl10, Ccl5*), implying diminished immune infiltration. Therefore, engineered PRC2-loss murine mammary tumor phenocopies the transcriptome changes and recapitulates the impaired immunogenicity of PRC2-loss human MPNST, irrespective of tumor lineage.

**Figure 6.**
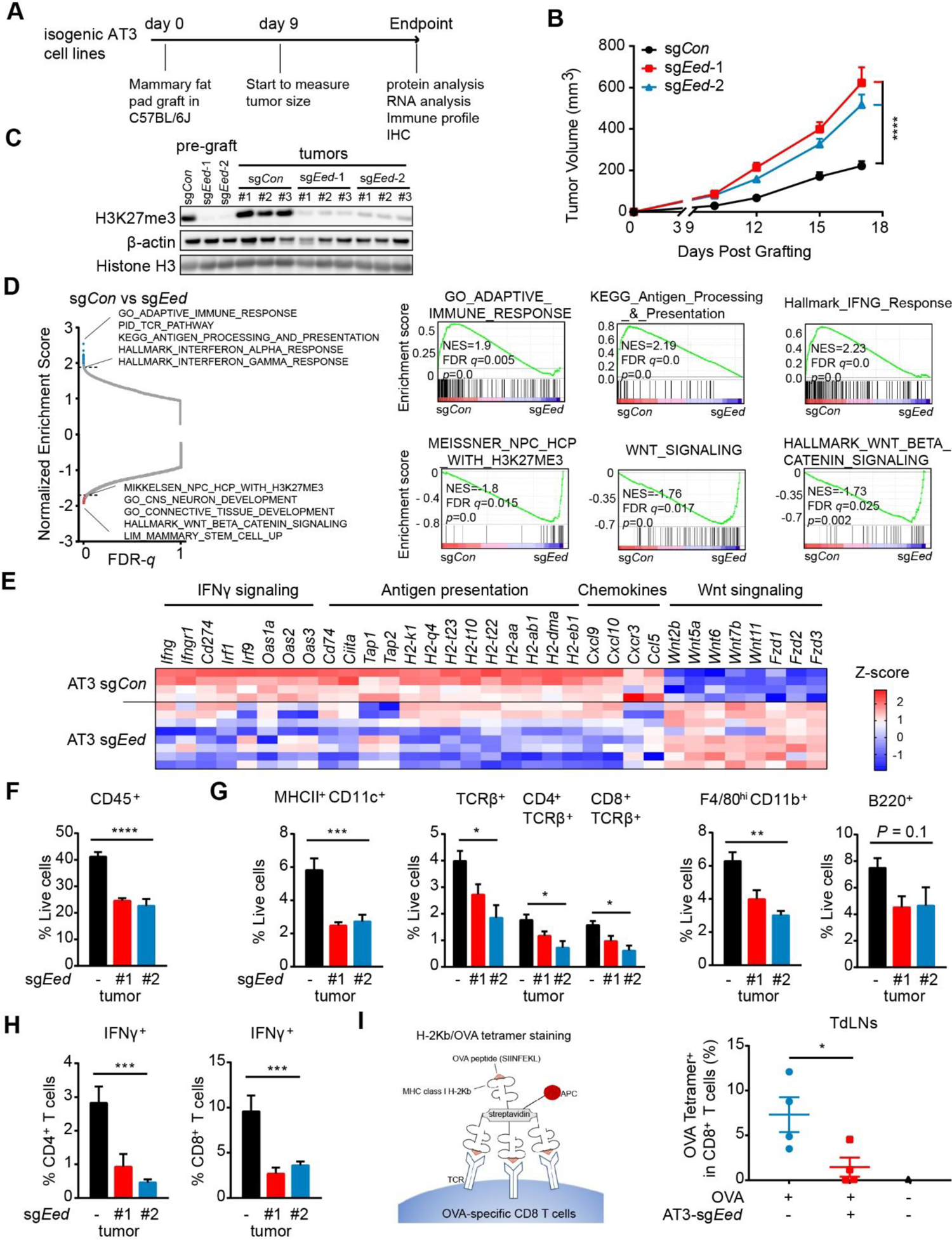
PRC2 loss leads to an antigen presentation defect and recapitulates the immune evasion phenotype in a murine mammary tumor syngeneic transplant model. **A**, A schematic of murine AT3 mammary tumor experimental plans *in vivo.* B, Representative growth curves of PRC2-isogeneic (sg*Con* vs. sg*Eed*) AT3 syngeneically transplanted tumors overtime. AT3 cells were orthotopically grafted bilaterally in the mammary fat pads in C57BL/6J mice. n=10 tumors in each group. **C**, Representative immunoblots of H3K27me3 in pre-graft AT3 cells and explanted AT3 tumor grafts. **D**, GSEA of transcriptomes derived from PRC2-isogenic AT3 (sg*Con* vs. sg*Eed*) syngeneically transplanted tumors. Left: GSEA normalized enrichment score (NES) vs. FDR *q*-value, which identified the most enriched gene sets with sg*Con* compared to sg*Eed*. Right: representative GSEA profiles of positively (upper) and negatively (bottom) enriched gene sets with PRC2-wt (sg*Con*) compared to PRC2-loss (sg*Eed*). sg*Con*: n=4; sg*Eed-*1: n=4; sg*Eed*-2: n=4; sg*Eed*-1 and sg*Eed*-2 transcriptomes are pooled together for analysis. **E**, Heatmap of expression changes for genes related to antigen presentation, IFNγ signaling response, chemokines and WNT signaling from RNA-seq of AT3 tumors. **F-G**, Percentage of CD45^+^ (F) and subpopulations (G), including DCs (B220^-^F4/80^lo^MHCII^+^CD11c^+^), T cells (TCRβ^+^), macrophages (B220^-^ F4/80^hi^CD11b^+^) and B cells (B220^+^), of tumor-associated immune infiltrates of total live cells in PRC2-isogenic AT3 syngeneically grafted tumors. n=5 for each group. **H**, The percentage of IFNγ^+^ cells of total CD4^+^ and CD8^+^ T cells in PRC2-isogenic AT3 tumors. n=10 for each group. \*\*\**P*<0.001 by unpaired two-tailed t test. **I**, Tetramer staining of OVA-specific CD8^+^ T cells in TdLNs 14 days after transplanting PRC2-isogenic AT3 cells in C57BL/6J mice. TdLNs of OVA^+^ tumors: n=4 for each group, TdLN of sg*Con* and OVA^-^ tumor: n=1. B and F-H: \**P*<0.05, \*\**P*<0.01, \*\*\**P*<0.001, \*\*\*\**P*<0.0001 by one-way ANOVA; I: \**P*<0.05 by unpaired two-tailed t test. All error bars: mean±SEM.

We next analyzed the population change of infiltrating immune cells in explanted PRC2-isogenic syngeneically transplanted tumors (**Fig. S5**). We observed significant reduction of CD45^+^ leukocytes in PRC2-loss tumors compared to PRC2-wt controls and confirmed by IHC (**Fig. 6F, S6D-S6E**). Similarly, the reduction of tumor immune infiltrates by PRC2 loss was across all major subclasses of immune cells, including MHCII^+^CD11c^+^ DCs, TCRβ^+^ T cells (both CD4^+^ and CD8^+^), F4/80^hi^CD11b^+^ macrophages, and to a lesser extent B220^+^ B cells (**Fig. 6G, S6F**). Importantly, functional IFNγ^+^ T cells were significantly reduced in both in CD4^+^ and CD8^+^ T cell subclasses (**Fig. 6H, S6G**), as well as a reduction of TNFα^+^ and a trend of reduction of the Granzyme B^+^ (GzmB^+^) CD8^+^ T cells in PRC2-loss tumor (**Fig. S6H-S6I**). These observations indicate that PRC2 loss in tumor not only lead to diminished numbers of T cells, but also significant functional impairment of T cells. Since functional T cells are the main source of IFNγ, these data indicate that IFNγ is likely significantly diminished in PRC2-loss tumors, which can further amplify the immune evasion phenotype.

To further dissect how PRC2 loss drive an immune evasion phenotype, we specifically evaluated the T cell priming in tumor draining lymph nodes (TdLNs), a critical initial step in the development of anti-tumor immunity [34, 41]. We exogenously expressed the model antigen, ovalbumin (OVA), in PRC2-isogenic AT3 cells (sg*Con* vs. sg*Eed*), and orthotopically transplanted these cells in the mammary fat pads of syngeneic C57BL/6J mice, and analyzed the OVA-specific CD8^+^ T cells in the TdLNs (**Fig. S6J**). The MHC-I OVA tetramer-positive CD8^+^ T cells were significantly diminished in the TdLNs from mice bearing PRC2-loss OVA^+^ AT3 tumors compared to PRC2-wt OVA^+^ ones (**Fig. 6I, S6K**). These data indicate that PRC2-loss in tumors suppresses the initial antigen cross-presentation by DCs and impairs the tumor-specific CD8^+^ T cell priming. These combined with the decreased tumor-infiltrating DCs and macrophages, as well as diminished immune recruitment chemokines, collectively contribute to the immune-desert TME. The observations from the AT3 murine mammary tumor model also demonstrate that the PRC2 loss-mediated immune evasion is not restricted to the MPNST context but can be generalized to other cancer contexts.

### Engineered PRC2-loss confers tumors the resistance to immune checkpoint blockades

Since tumor cell-intrinsic PRC2 loss drives an immune-desert TME, we speculated that PRC2 loss might confer primary resistance to FDA-approved immune checkpoint blockade immunotherapies, including anti-CTLA4 and anti-PD1 antibodies [42]. We evaluated the therapeutic efficacy of combining anti-CTLA4 and anti-PD1 antibodies in transplanted PRC2-isogenic AT3 tumor models (**Fig. 7A**). Combined ICB was effective and significantly inhibited the growth of PRC2-wt AT3 tumors. By contrast, it failed to retard the growth of PRC2-loss AT3 tumors (**Fig. 7B**). Along with tumor growth inhibition, the combined ICB treatment increased CD45^+^ immune infiltrates (**Fig. 7C-7D, S7A**), particularly TCRβ^+^CD4^+^ and TCRβ^+^CD8^+^ T cell infiltrates in the PRC2-wt tumors (**Fig. 7E-7F, S7B-S7C**). However, the recruitment of CD45^+^ immune cells, including both CD4^+^ and CD8^+^ T cells, was significantly blunted in the PRC2-loss tumors. The functional IFNγ^+^ CD4 and CD8 T cells trended to reduce in PRC2-loss tumors compared to PRC2-wt tumors in response to combined ICB treatment (**Fig. 7G**). These results support that PRC2-loss tumors are resistant to ICB therapy.

**Figure 7.**
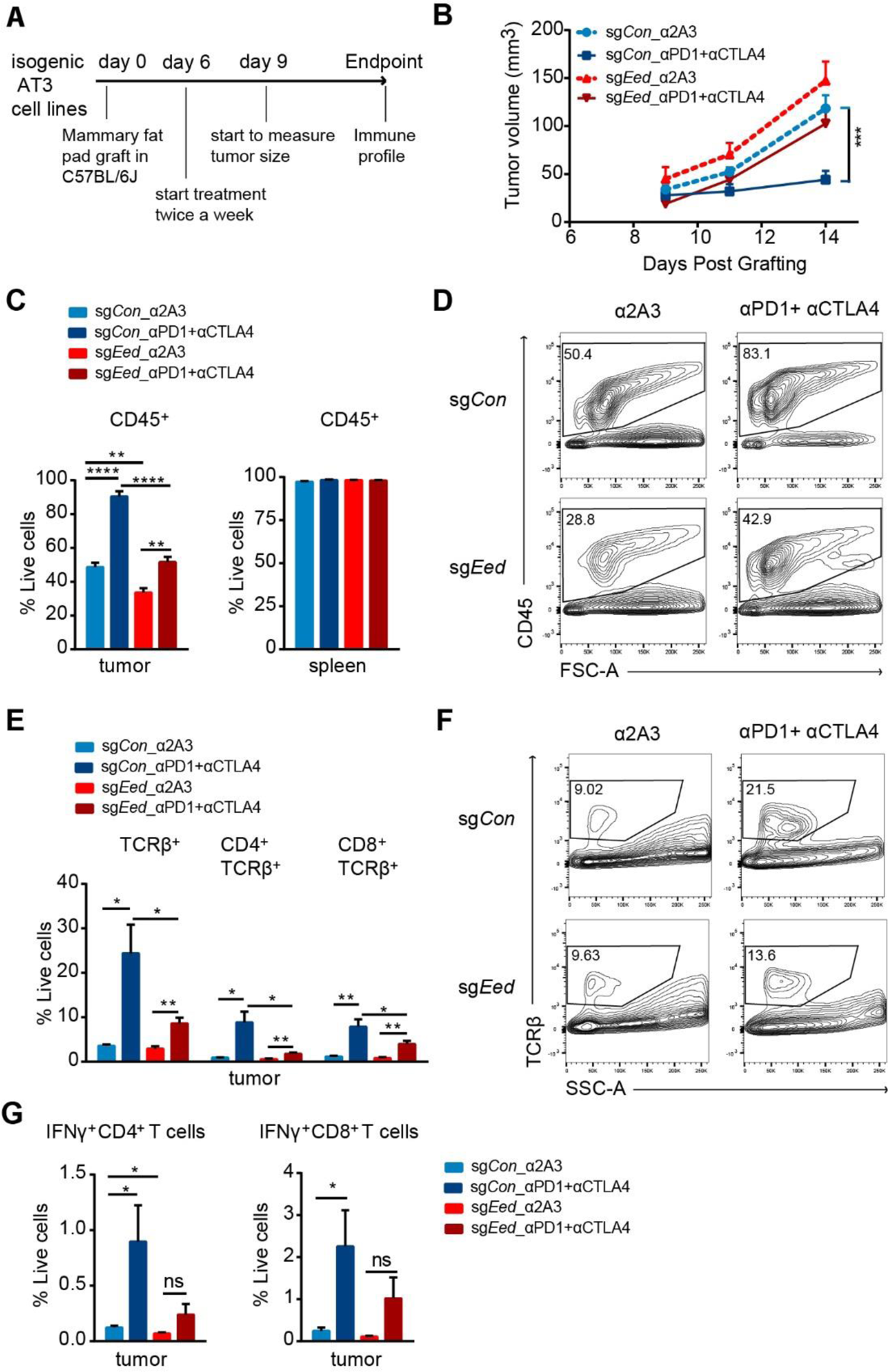
PRC2-loss tumors confer primary resistance to immune checkpoint blockade (ICB) therapies. **A**, A schematic of treatment plan of PRC2-loss AT3 tumors with ICB. **B**, Tumor growth curves of PRC2-isogenic AT3 tumors with ICB treatment overtime. PRC2-isogenic AT3 cells (sg*Con* vs. sg*Eed*) were orthotopically grafted bilaterally in the mammary fat pads of C57BL/6J mice, and mice were treated with intraperitoneal injection of 250 µg anti-PD1 and 200 µg anti-CTLA4 combination or 100 µg isotype-matched anti-2A3 control antibody. n=10 tumors for each group. **C**, Percentage of CD45^+^ tumor immune cells in PRC2-isogneic AT3 tumors (left) or spleens (right). n=5 for each cohort. **D**, Representative examples of CD45^+^ cells (gated CD45^+^ cells in live cells) by FACS. **E**, Percentage of T cells (TCRβ^+^), CD4^+^ and CD8^+^ subpopulation of T cells in PRC2-isogenic AT3 tumors treated with ICB and controls. n=5 for each cohort. **F**, Representative examples of T cells (gated TCRβ^+^ in CD45^+^ cells) by FACS. **G**, Percentage of IFNγ^+^ CD4^+^ and CD8+ T cells in all live cells in tumors. n=5 for each cohort. B, C, E and G: \**P*<0.05, \*\**P*<0.01, \*\*\*\**P*<0.0001 by unpaired two-tailed t test. All error bars: mean±SEM.

### Intratumoral delivery of immunostimulatory heat-inactivated MVA sensitizes PRC2-loss tumors to ICB therapy

We hypothesized that therapeutic strategies that can enhance the interferon response and innate immunity might modulate the immune-desert TME of PRC2-loss tumors. To test that, we investigated whether intratumoral (IT) delivery of inactivated modified vaccinia virus Ankara (MVA) sensitize the PRC2-loss tumors to ICB. MVA infection of DCs induces type I IFN via activating the cytosolic DNA-sensing pathway mediated by cyclic GMP-AMP synthase (cGAS) and Stimulator of IFN genes (STING) [43]. Previous studies have shown that heat-inactivation prevents the production of viral inhibitory proteins, and heat-inactivated MVA (heat-iMVA) induces higher levels of type I IFN than live MVA [44]. In addition, IT heat-iMVA generates local and systemic anti-tumor immunity, which requires CD8^+^ T cells and Batf3-dependent CD103^+^/CD8^+^ DCs, and is dependent on STING [44]. Here, we evaluated whether IT heat-iMVA could disrupt the “cold” TME and sensitize PRC2-loss tumors to combined ICBs (anti-PD1 plus anti-CTLA4) using the engineered ICB-resistant PRC2-loss AT3 (sg*Eed*) in the syngeneic transplant murine model system (**Fig. 8A**). IT delivery of heat-iMVA alone only mildly retarded the tumor growth of PRC2-loss AT3 (sg*Eed)* tumor, whereas IT heat-iMVA significantly reduced tumor growth when combined with anti-PD1 and anti-CTLA4 (**Fig. 8B, S8A**), accompanied by significant prolongation of survival of the heat-iMVA and ICB combination treatment group compared to ICB treatment alone or vehicle groups (**Fig. 8C**). We also observed significantly more cell death in PRC2-loss tumors under the triple combination therapy (**Fig. S8B**), suggesting an enhanced antitumor effect. Consistently, IT injection of heat-iMVA significantly increased CD45^+^ immune cell infiltration in PRC2-loss AT3 (sg*Eed*) tumors compared to vehicle, further more when combined with ICBs (**Fig. 8D**), including TCRβ^+^ T cells and MHCII^+^CD11c^+^ DCs (**Fig. 8E-8F, S8C-S8D**); no significant changes in F4/80^hi^CD11b^+^ macrophages and B220^+^ B cells were observed in heat-iMVA injected tumors (**Fig. S8E-S8F**). Both CD4^+^ and CD8^+^ T cells were significantly enriched accompanied by increased proliferation (Ki67^+^) in heat-iMVA treatment group with and without combination with ICB, compared to vehicles (**Fig. 8E-8F**). Moreover, heat-iMVA treatment led to significant decrease in immune suppressive FoxP3^+^ Treg cells (**Fig. 8G**), and significant increase in GzmB^+^CD8^+^ cytotoxic T cells compared to vehicles (**Fig. 8H**), especially when combined with ICB therapy. These results imply that IT delivery of MVA-based viral therapy combined with ICB may be an effective initial strategy to modify the cold TME and elicit anti-tumor effect in PRC2-loss tumors.

**Figure 8.**
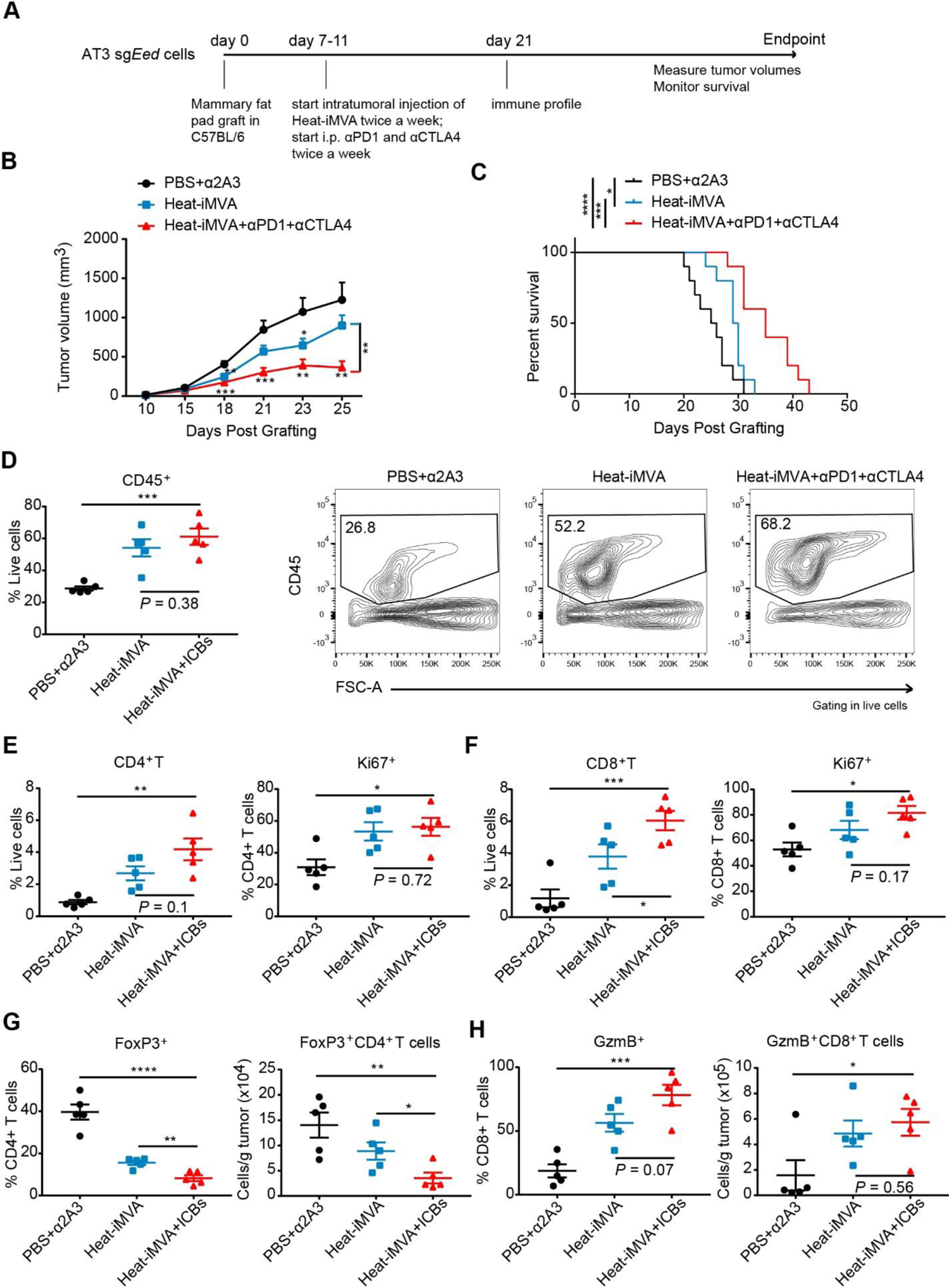
Intratumoral injection of heat-iMVA sensitizes PRC2-loss tumors to anti-PD1 and anti-CTLA4 therapy in engineered AT3 murine breast cancer model. **A**, A schematic of treatment plan of heat-iMVA and ICBs in PRC2-loss AT3 (sg*Eed*) tumors. **B-C**, Tumor growth curves (B) and Kaplan-Meier survival curve (C) overtime in mice with AT3 sg*Eed* tumors with indicated treatments. AT3 sg*Eed* cells were orthotopically grafted in the right mammary fat pads of C57BL/6J mice. Dose of treatments: an equivalent of 8×10^7^ plaque-forming units (pfu) heat-iMVA, 250 µg anti-PD1, and 200 µg anti-CTLA4, or combination per tumor. n=10 tumors for each group. **D**, Percentage (left) and representative FACS profiles (right) of tumor infiltrating CD45^+^ immune cells in AT3 (sg*Eed*) tumors under different treatment conditions as indicated. n=5 for each cohort. **E-F**, Percentage of CD4^+^ T cells (E) and CD8^+^ T cells (F) in all live cells and the relative Ki67^+^ subpopulation in AT3(sg*Eed*) tumors under various treatment conditions as indicated. **G-H**, Percentage (left) and absolute number (right) of FoxP3^+^ Treg cells in CD4^+^ T cells G) and GzmB^+^ cells in CD8^+^ T cells (H) in AT3 (sg*Eed*) tumors under various treatment conditions as indicated. n=5 for each cohort. ICBs: anti-PD1 + anti-CTLA4. \**P*<0.05, \*\**P*<0.01, \*\*\**P*<0.001, \*\*\*\**P*<0.0001, B: by unpaired two-tailed t test; C: by Log-rank (Mantel-Cox) test; D-H: by one-way ANOVA among all three groups, and by unpaired two-tailed t test between heat-iMVA and heat-iMVA + ICB. All error bars: mean±SEM.

## Discussion

Unlike previously described mechanisms of tumor cell-intrinsic immune evasion that primarily affect selective subpopulations of tumor immune infiltrates, e.g., decreased T lymphocyte infiltration and function, or recruitment of myeloid suppressive cells (e.g., *LKB1* mutation in lung cancer) [41, 45–53], PRC2 loss in tumor affects a broad spectrum of subpopulations of immune infiltrates, including dendritic cells, T cells, B cells, and/or macrophages, and therefore, drives an immune-desert TME. Our data indicate that PRC2 loss impacts multiple anti-tumor immune response pathways, including impaired antigen presentation and T cell priming, reduced chemotactic cytokine production (e.g., *CCL2*, *CCL5*) resulting in diminished innate and adaptive immune cell recruitment, and dampened IFNγ response.

Mechanistically, PRC2 loss in tumor cells leads to genome-wide redistribution of the chromatin accessibility and chromatin modifications (e.g., H3K27ac), and hence reprograming of the chromatin landscape. Although with PRC2 loss, global H3K27me2/3 histone modifications are lost with a reciprocal global gain of H3K27ac on chromatin, the genome-wide distribution of gained H3K27ac is not uniform. The increase of H3K27ac is not due to the activity of the canonical histone acetyl transferase (HAT) for H3K27ac, CBP/P300, as other classical CBP/P300 substrates, e.g., H3K9ac and H3K18ac, have remained stable with PRC2 loss (data not shown). We speculate that the global increase of H3K27ac level is due to increased availability of H3K27 substrate as a result of H3K27me1/2/3 loss in PRC2-loss tumors. Consistent with chromatin accessibility changes mediated by PRC2 loss, there are many genes with significantly increased H3K27ac enrichment at distal regulatory enhancers and intergenic regions with corresponding increase of transcriptomes; these included master regulators, developmental and lineage specification pathways, and importantly genes involved in the WNT signaling pathway that has previously been shown to actively suppress T cell recruitment and affect the local anti-tumor response through defects in T cell priming [41, 51, 52, 54]. Curiously, PRC2 loss also led to a paradoxical decrease of H3K27ac enrichment at many genome-wide loci, mainly at active promoters and distal regulatory enhancers including existing super enhancers in PRC2-wt tumor cells, and corresponding decrease in transcriptomes. This negative impact of H3K27ac enrichment preferentially affected the immune signaling pathways, including chemotactic cytokines (e.g., *CCL2*), type I and type II IFN response genes and others, leading to significantly diminished expression of chemokines and dampened IFNγ signaling responses *in vitro,* and the diminished anti-tumor immune response was further amplified *in vivo*. Thus, in the relevant cell context, PRC2 loss reprograms the chromatin landscape and shifts a baseline primed immune signaling-dependent cellular response to the PRC2-regulated development and cellular differentiation transcriptional programs. Through these mechanisms, PRC2 loss reprograms the tumor cell and the TME to decrease antigen presentation and reduce chemotactic cytokine secretion, leading to diminished tumor immune infiltrates and ICB primary resistance. Our observations are also consistent with recent reports demonstrating that inactivating mutations of the core components (e.g., *PBRM1*) of the PBAF complex that are antagonistic to PRC2 complex in tumor cells enhance IFNγ signaling and sensitize the PBAF-deficient tumors to ICB therapies [55, 56].

Viral-based cancer immunotherapy is a promising strategy to induce host anti-tumor immunity through multiple mechanisms, including viral-induced oncolysis and alteration of immunosuppressive TME [57, 58]. Modified vaccinia virus Ankara (MVA) is a highly attenuated vaccinia strain recently approved as a safe and effective vaccine against smallpox and monkeypox and has been investigated as a promising vaccine vector [59, 60]. Infection of tumor and immune cells by IT delivery of heat-iMVA leads to robust induction of type I IFN and proinflammatory cytokines and chemokines to attract and activate immune effector cells [44]. Here, we showed that IT heat-iMVA disrupted immune tolerance and activated immune cells to eliminate malignancies in PRC2-loss tumors, particularly when combined with other immunotherapies. Furthermore, engineered recombinant MVA by deleting immunosuppressive viral genes, and expressing T cell-activating cytokines to enhance immune response and anti-tumor activity, are currently being developed as next-generation of novel viral immunotherapies, which can be evaluated clinically in PRC2-loss tumors.

Genetic and functional inactivation of PRC2 is prevalent in MPNST and a variety of other cancer types. Our data point to PRC2-loss as a novel tumor cell-intrinsic mechanism of an immune desert TME and primary resistance to ICB. PRC2 loss can be readily identified by loss of immunostaining of H3K27me3 in tumor specimens clinically [3, 13, 14]; therefore, H3K27me3 immunostaining can be immediately utilized as a molecular biomarker to identify clinical cases that are unlikely to benefit from ICB therapy and to salvage patients from ICB treatment-related toxicity. Moreover, our studies have identified therapeutic viral therapy that enhances tumor immunity as an initial strategy to overcome the immune-desert TME of PRC2-loss tumors and capitalize on ICB therapy.

## Methods

### Human tumor tissue collection

Clinical samples were collected during surgical resection from patients diagnosed as MPNST according to Memorial Sloan Kettering Cancer Center (MSKCC) Institutional Review Board (IRB) protocol. All patients provided informed consent. Frozen and paraffin embedded tissue samples were banked, and tissue microarrays (TMAs) were generated. All MPNST were pathologically reviewed and confirmed by a sarcoma expert (C.R.A.) at MSKCC. Sample annotation is shown in Supplementary Table 1.

### Cell lines

HEK-293T cell line was purchased from ATCC. The human MPNST cell line M3 (NF1^-/-^, CDKN2A/B^-/-^) was a gift and developed from NF1-associated MPNT patient by William Gerald. The mouse MPNST cell line SKP605 (*Nf1*^-/-^, *Cdkn2a/b*^-/-^) was generated from skin-derived precursors by A.J.P. in our lab according to the published protocol [40] and confirmed with pathologist (C.R.A.). (**Fig. S4A-S4B**). The mouse breast cancer cell line AT3 was a gift from the laboratory of M.L. at MSKCC. All cells were tested to be mycoplasma free by the MycoAlertPlus MycoPlasma Detection Kit (Lonza). All these cells were cultured in DMEM supplemented with L-glutamine (2 mM), penicillin (100 U/ml), and streptomycin (100 μg/ml), and 10% heat-inactivated fetal bovine serum (FBS) in 5% CO2 incubator at 37°C.

### Immunohistochemistry (IHC)

Human tissue processing, embedding, sectioning and staining with hematoxylin and eosin were performed by the MSKCC Department of Pathology. IHC of human TMA tumor samples were performed by the MSKCC Human Oncology and Pathogenesis Program automatic staining facility using a Ventana BenchMark ULTRA Automated Stainer. Mouse fresh tissues were fixed with 4% PFA overnight, washed with PBS 3 time, 5min for each time, and stored in 70% ethanol. Tissue paraffin embedding, sectioning, and H&E staining were performed by the Histoserv, Inc. IHC was performed by the Molecular Cytology Core Facility at MSKCC for immune marker CD45, or Ventana BenchMark ULTRA Automated Stainer for H3K27me3 (Millipore #07-449, 1:500, protocol#313), Ki67 (Abcam #ab15580, 1:500, protocol#312) and S100B (Abcam #ab52642, 1:2000, protocol#313). Slides were scanned by the Molecular Cytology Core Facility at MSKCC and analyzed using CaseViewer software.

### Gene knockout by CRISPR/Cas9

pLCP2B plasmid was generated by removing Cas9-P2A-tRFP from pL-CRISPR.EFS.tRFP (Addgene, #57819) and replacing with Cas9-P2A-Blast in our lab. lentiCRISPR-v2 vector with puromycin was purchased from Addgene #52961. The sgRNA oligos as below were annealed, digested using *BsmB*I (New England BioLab, #R0580L) and cloned into vectors, such as pLCP2B or lentiCRISPR-v2. To make lentivirus, the transfer plasmid was co-transfected into HEK-293T cells with the packaging plasmids pVSVg (Addgene, #8454) and psPAX2 (Addgene, #12260). To delete target genes, cells were transduced with respective sgRNA and selected with 5 - 10 μg/ml blasticidin S HCl (Thermo Fisher, # A1113903) or 2 - 5 μg/ml Puromycin Dihydrochloride (Thermo Fisher, # A1113803) until all negative control cells were dead. Then M3 sg*Con* were M3 sg*SUZ12* cells were seeded at super-low density to allow colony formation from single cells in 96-well plates. Colonies were then picked and expanded for knockout validation by immunoblotting based on H3K27me3 level. Similarly, sg*Nf1* and sg*Cdkn2a* were cloned into the pLCP2B and blasticidin S HCl selected; sg*Cdkn2b* was cloned in LentiCRISPRv2-mCherry vector (Addgene, #99154) in SKP cells. sg*Eed* was were cloned into lentiCRISPR-v2, and SKP605 cells were puromycin selected and single clone picked. M3 sg*Con* cells were pooled from 6 single clones, while M3 sg*SUZ12* cells were pooled from 9 single clones. Both SKP605 sg*Con* cells and sg*Eed* cells were 4 single clones pooled. The doubling time for M3 cells is about 48 hours, while the doubling time for SKP605 is about 24 hr. The cells will be used for experiments within 1 month (15-30 passages) after pooling single clones.

sgRNA Oligos are as following Sequence (5’ to 3’) and their PAM are NGG:

sg*Con* (Control): F, caccgGCTGATCTATCGCGGTCGTC, R, aaacGACGACCGCGATAGATCAGCc; Human_sg*SUZ12*_VEFS: F, caccgTCCATTTCTTGTGGACGGAG, R, aaacCTCCGTCCACAAGAAATGGAc; Mouse_sg*E*ed-1_WD40: F, accgGAAGGTTTGGGTCTCGTGGG, R, aaacCCCACGAGACCCAAACCTTCc; Mouse_sg*Eed*-2_WD40: F, caccgGGAGAAGGTTTGGGTCTCGT, R, aaacACGAGACCCAAACCTTCTCCc; Mouse_sg*Suz12*_VEFS:F, tcccGAATTTTCTGATGTGAATGA, R, aaacTCATTCACATCAGAAAATTC; Mouse_*Nf1*: F, caccgGGGAGAACTCCCTATAGCTA, R, aaacTAGCTATAGGGAGTTCTCCCc; Mouse_sg*Cdkn2a*: F, caccgCGGTGCAGATTCGAACTGCG, R, aaacCGCAGTTCGAATCTGCACCGc; Mouse_sg*Cdkn2b*: F, caccgTTGGGCGGCAGCAGTGACGC, R, aaacGCGTCACTGCTGCCGCCCAAc;

### Transplant mouse model

Wild-type 6-8-week-old female C57BL/6J mice were purchased from the Jackson Laboratory (Stock No: 000664), while 6-8-week-old female NSG mice were purchased from MSKCC core facility. All the procedures related to mouse handling, care and the treatment were performed according the guidelines from the MSKCC approved by the Institutional Animal Care and Use Committee (IACUC). Animal protocols were approved by the MSKCC. Isogenic M3 cells were orthotopically transplanted into sciatic nerve pockets of NSG mice: 3 million cells in 100 μl precooling 1:1 PBS:Matrigel (Corning, #356237). Isogenic SKP605 cells were orthotopically transplanted into sciatic nerve pockets of C57BL/6J mice: 5 million cells in 100 μl precooling 1:1 PBS:Matrigel. Isogenic AT3 cells were orthotopically transplanted into mammary fat pads of C57BL/6J mice: 100 to 150 thousand cells in 100 μl precooling PBS. Staples were removed at least after 1 week of grafting. Tumors were measured twice a week by Vernier caliper. Tumor volume (TV) = 4/3*pi*length/2*width/2*height/2. For survival study, mice were euthanized when the tumor volume reached 1500 mm^3^ or the mice exhibit signs of illness.

### Monoclonal antibody therapy

Monoclonal antibodies anti-PD1, anti-CTLA4 and anti-2A3 were purchased from Bio X cell: Anti-mouse PD-1 (CD279) (#BE0146), anti-mouse CTLA-4 (CD152) (#BE0164), and rat IgG2a isotype control, anti-trinitrophenol (#BE0089). Antibodies were i.p. once every three days at a dose of 250 μg anti-PD1 + 200 μg anti-CTLA4 or 100 μg anti-2A3 per 100 μl PBS per mouse per treatment.

### Viruses and intratumoral injection with viruses

The MVA virus was kindly provided by G. Sutter (University of Munich) to the Deng laboratory. MVA was propagated in BHK-21 cells (baby hamster kidney cell, American Type Culture Collection (ATCC) CCL-10) and purified through a 36% sucrose cushion. Heat-iMVA was generated by incubating purified MVA virus at 55° C for 1 hour (Dai et al., Science Immunology 2017). At 7 to 11 days after implantation, tumors were measured and intratumorally injected with heat-iMVA (an equivalent of 4 x 10^8^ plaque-forming units) in 200 μl or PBS twice weekly when mice were under anesthesia. Mice were monitored daily.

### Tumor infiltrate analysis by flow cytometry

Fresh tumor tissues were minced with razor blades on slides one by one, then digested with 10 ml digestion buffer separately (MPNST tumors: 200 U/ml Collagenase I + 300 U/ml Collagenase IV + 4 μg/ml DNase I + RPMI-1640, breast cancer tumors: 300 U/ml Collagenase III + 4 μg/ml DNase I + HBSS). All these enzymes were purchased from Worthington: Collagenase I #LS004196, Collagenase III #LS004182, Collagenase IV # LS004188, and DNase I #LS002006. HBSS buffer was purchased from Thermo Fisher Scientific #14025076. Samples were incubated for 1 hr in 37°C shaker. To get single cells, tumors were mashed through 70micron filters. To enrich live cells, Ficoll (Thermo Fisher Scientific, #17144002) centrifugation was used with subsequent washing of the obtained cells. Spleens were minced directly and lysed in ACK lysis buffer (Thermo Fisher Scientific, #A1049201) to remove red cells. For flow cytometric analysis, about 1.5 million washed cells were resuspended in 70μl FACS buffer (2% FBS + 1 mM EDTA (0.5M, 1:500) + 10 mM HEPES (1M, 1:100) in PBS) with Fc block for 15 min on ice first, then Fc block plus indicated antibodies for 30 min on ice from light. Foxp3 / Transcription Factor Staining Buffer Kit (Tonbo, #TNB-0607-KIT) was used to stain intracellular markers. All FACS were performed on BD LSR Fortessa, and data were analyzed using FlowJo software. Antibodies for FACS are as below: Purified Anti-Mouse CD16 / CD32 (1:500, 2.4G2) (TONBO, #70-0161-U100), Ghost Dye™ Violet 450 (1:1000, TONBO, #13-0863-T100), PE-Texas Red_CD45 (1:200, Thermo Fisher Scientific, #MCD4517), APC/Cy7_TCR β (1:200, Biolegend, #109220), PerCP-Cyanine5.5_ CD8a (1:200, TONBO, #65-1886-U100), BV510_CD4 (1:200, BD, #563106), PE_B220 (1:400, TONBO, #50-0452-U100), FITC_F4/80 (1:200, TONBO, #35-4801-U100), APC-Cyanine7_CD11b (1:200, TONBO, #25-0112-U100), PE-Cyanine7_CD11c (1:200, TONBO, #60-0114-U025), APC_MHC Class II (I-A/I-E) (1:600, TONBO, #20-5321-U100), FITC_IFN gamma (1:100, TONBO, #35-7311-U100), PE-Cyanine7_TNF alpha (1:100, Thermo Fisher Scientific, #25-7423-82)

### T-cell stimulation

Cell Stimulation Cocktail (plus protein transport inhibitors) (500X) (Thermo Fisher, #00-4975-93) was diluted in T cell medium (RPMI 1640 + 10%FBS + L-glutamine (2 mM) + penicillin (100 U/ml + streptomycin (100 μg/ml) + 55 μM 2-Mercaptoethanol (Thermo Fisher, #21985023)). 2 million cells were suspended in 200 μl T-cell medium with stimulation cocktail in plates, then incubated for 4 hours at 37° incubator. After incubation, cells were harvested, stained with surface markers and then intracellular markers for FACS analysis.

### OVA model antigen system

pMSCV-EGFP-PGK-Luc2-2A-USA plasmid was a gift from Dr. Ming Li laboratory at MSKCC, which include OT-I binding antigen and OT-II binding antigen for MHC of C57BL/6J mouse background. To make retrovirus, HEK293T cells were transfected with pMSCV-EGFP-PGK-Luc2-2A-USA, packaging plasmids pVSVg (Addgene, #8454) and pEco (Takara, #PT3749-5). Cells transduced with the retrovirus were sorted for EGFP positive by flow cytometry, and these cells stably overexpressed OVA model antigens: OT-I binding antigen and OT-II binding antigen. APC anti-mouse H-2Kb bound to SIINFEKL antibody (BioLegend, #141605) was used to detect OVA model antigen on cancer cell surface. To detect OVA-specific T cell priming, OVA+ and OVA-AT3 cells were orthotopically grafted in mammary fat pads of C57BL/6 mice. After 18 days post grafting, cells isolated from tumor draining lymph nodes (TdLNs) were incubated with 2μg/ml OVA 257-264 in T cell medium for 24 h at 37° incubator. First, cells were stained with Fc blocking for 15 mins; Second, Fc blocking plus iTAg H-2Kb OVA Tetramer-SIINFEKL-APC (MBL, #TB-5001-2) in FACS buffer for 1h at room temperature from light. Third, live/dead dye (TONBO, #13-0863-T100) and anti-mouse CD8a-FITC (TONBO, #35-1886-U100) were added into the staining system for 30 mins at 4°. Finally, cells were washed by FACS buffer for 3 times and ready for FACS analysis.

### Cell colony formation assay

AT3 cells were seeded into 6-well plates at a concentration of 200 cells per well. After 9 days. colonies were fixed with fixation solution (10% methanol + 10% acetic acid) at room temperature for 15 min and then stained with a solution of 1% crystal violet in methanol for 15 min.

### Protein extraction and western blotting

After washed by precooling PBS, cells were lysed using preheated 2% SDS followed by boiling for 30 min at 95°. Protein concentrations were determined using BCA protein assay kit (Thermo Fisher, #23225). Protein samples were prepared by adding LDS loading buffer (4X) (Thermo Fisher, #NP0008) and 1 M DTT followed by boiling for 30 min at 95°. Proteins were separated by 4-12% Bis-Tris Gel (Thermo Fisher, #NP0336BOX) with MES SDS running buffer (20X) (Thermo Fisher, #NP0002), and transferred to the nitrocellulose membrane (BioRad, #1620115). Membranes were blocked with blocking buffer (Thermo Fisher, #UH289384) for 1 hr at room temperature and incubated with the indicated primary antibodies at 4° overnight. After washing with 1xTris-buffered saline Tween-20 (TBST) (25 mM Tris, 150 mM NaCl, 2 mM KCl, pH 7.4, supplemented with 0.2% Tween-20) three times for 30 min, membranes were incubated with peroxidase-conjugated secondary antibodies at room temperature for 1 hr. The membranes were washed again with TBST three times and visualized with chemiluminescence using HRP substrate (Millipore, #WBKLS0500) or Pico (Thermo Fisher, #34578).

Primary antibodies for immunoblotting are as below: anti-NF1 (1:2000, Bethyl, #A300-140A), anti-CDKN2A (1:1000, DeltaBioLabs, #DB018), anti-CDKN2B (1:500, Abcam, #ab53034), anti-SUZ12 (1:1000, CST, #3737), anti-H3K27me3 (1:2000, CST, #9733), anti-H3K27me2 (1:5000, CST, #9728), anti-H3K27me1 (1:1000, Takara, #MABI0321-100I), anti-H3K27ac (1:4000, Abcam, #ab4729), anti-Histone H3 (1:2000, CST, #12648), anti-β-actin (1:5000, Proteintech, #66009-1-Ig), anti-Lamin B1 (1:2000, Proteintech, #12987-1-AP) and anti-GAPDH (1:3000, CST, #5174S).

### RNA isolation and qRT-PCR

Total RNA was isolated from cell lines using total RNA kit I (Omega, #R6834-02) and homogenizer mini columns (Omega, #HCR003), or tissues using Trizol (Thermo Fisher #15596026). cDNA was prepared using High-Capacity cDNA Reverse Transcription Kit (Thermo Fisher, # 4368814). qRT-PCR was performed following the instruction for SYBR Green Master Mix (Thermo Fisher, #A25777) with V7 Real-Time PCR system (Applied Biosystems). Expressed values relative to control were calculated using the ΔΔCT method. Housekeeping gene RPL27 was used as reference gene for normalization. The sequences of primers used for qPCR analysis were listed as follows (5’--3’).

Human_*RPL27*: F, CATGGGCAAGAAGAAGATCG, R TCCAAGGGGATATCCACAGA; Human_*SUZ12*: F, TTGCAGCTTACGTTTACTGGTT, R, GGAACTTGCCTTATTGGACAACT; Human_*EED*: F, CTGTAGGAAGCAACAGAGTTACC, R, CATAGGTCCATGCACAAGTGT; Human_*CD274*: F, TGGCATTTGCTGAACGCATTT, R, TGCAGCCAGGTCTAATTGTTTT; Human_*IRF1*: F, CTGTGCGAGTGTACCGGATG, R, ATCCCCACATGACTTCCTCTT; Human_*CCL2*: F, CAGCCAGATGCAATCAATGCC, R, TGGAATCCTGAACCCACTTCT; Human_*CD74*: F, GATGACCAGCGCGACCTTATC, R, GTGACTGTCAGTTTGTCCAGC; Human_*CIITA*: F, CCTGGAGCTTCTTAACAGCGA, R, TGTGTCGGGTTCTGAGTAGAG; Human_*HLA-DRA*: F, AGTCCCTGTGCTAGGATTTTTCA, R, ACATAAACTCGCCTGATTGGTC; Human_*HLA-DRB*: F, CGGGGTTGGTGAGAGCTTC, R, AACCACCTGACTTCAATGCTG; Human_*LAP3*: F, GTCTGGCCGTGAGACGTTT, R, ACCATAAAAGGTTCGAGTCTTCC; Mouse_*Rpl27*: F, AAAGCCGTCATCGTGAAGAAC, R, GATAGCGGTCAATTCCAGCCA; Mouse_*Eed*: F, AGCCACCCTCTATTAGCAGTT, R, GCCACAAGAGTGTCTGTTTGGA; Mouse_*Ccl2*: F, TAAAAACCTGGATCGGAACCAAA, R, GCATTAGCTTCAGATTTACGGGT; Mouse_*Cxcl10*: F, CCAAGTGCTGCCGTCATTTTC, R, TCCCTATGGCCCTCATTCTCA; Mouse_*Il2*: F, AACCTGAAACTCCCCAGGAT, R, TCATCGAATTGGCACTCAAA; Mouse_*Oas1*: F, GGGCCTCTAAAGGGGTCAAG, R, TCAAACTTCACTCCACAACGTC; Mouse_*Cd274*: F, AGTATGGCAGCAACGTCACG, R, TCCTTTTCCCAGTACACCACTA; Mouse_*Irf1*: F, GGCCGATACAAAGCAGGAGAA, R, GGAGTTCATGGCACAACGGA; Mouse_*Actb*: F, GTGACGTTGACATCCGTAAAGA, R, GCCGGACTCATCGTACTCC

### RNA-seq and analysis

Total RNA was isolated from fresh tissues or duplicate wells of indicated conditions using Trizol (Thermo Fisher #15596026). RNA-seq library construction and sequencing were performed at the integrated genomics operation (IGO) core facility at MSKCC using poly-A capture. The libraries were sequenced on an Illumina HiSeq-2500 platform with 50-bp paired-end (human MPNST tumors) or single-end (AT3 tumors) reads to obtain a minimum yield of 40 million reads per sample. The sequence data were processed and mapped to the human reference genome (hg19) or mouse reference genome (mm9) using STAR v2.3 [61]. Gene expression was quantified to transcripts-per-million (TPM) using the STAR [62] and Log 2 transformed. GSEA was performed using JAVA GSEA 2.0 program [63].

### Assay for transposase-accessible chromatin using sequencing (ATAC-seq) and analysis

Cells were washed with PBS, trypsinized from 6-well plates and harvested as a single-cell suspension in precool PBS. 100 thousand cells were sent to Center for Epigenetics Research at MSKCC for ATAC-seq performed as previously described [64] [65]. For each sample, cell nuclei were prepared from 50,000 cells, and incubated with 2.5 μl of transposase (Illumina) in a 50-μl reaction for 30 min at 37 °C. After purification of transposase-fragmented DNA, the library was amplified by PCR and subjected to paired-end 50 base-pair high-throughput sequencing on an Illumina HiSeq2500 platform.

For data analysis, ATAC-seq reads were quality and adapter trimmed using Trim Galore before aligning to human genome assembly hg19 with Bowtie2 using the default parameters. Aligned reads with the same start position and orientation were collapsed to a single read before subsequent analysis. Density profiles were created using the BEDTools suite, with subsequent normalization to a sequencing depth of ten million reads for each library. To ascertain regions of open chromatin, MACS2 (https://github.com/taoliu/MACS) was used with a p-value setting of 0.001 against a cell line-matched input sample. A global peak atlas was created by first removing blacklisted regions then merging all peaks within 500 bp and counting reads with version 1.6.1 of featureCounts (http://subread.sourceforge.net). Differential enrichment was scored using DESeq2 for all pairwise group contrasts. Differential peaks were then merged into a union set, and k-means clustering was performed from k=4:10, stopping when redundant clusters emerged. Peak-gene associations were created by assigning all intragenic peaks to that gene, and otherwise using linear genomic distance to transcription start site. GSEA (http://software.broadinstitute.org/gsea) was performed with the pre-ranked option and default parameters, where each gene was assigned the single peak with the largest (in magnitude) log2 fold change associated with it. Motif signatures were obtained using HOMER v4.5 (http://homer.ucsd.edu).

### Chromatin Immunoprecipitation sequencing (ChIP-seq) and analysis

Chromatin isolation from indicated cells and immunoprecipitation were performed as previously described [66]. The libraries were sequenced on an Illumina HiSeq2500 platform with 50-bp paired-end reads. Reads were trimmed by the software Trim Galore and then aligned to the human genome (hg19) using the Bowtie2 alignment software (v2.3.5) [67]. Duplicated reads were eliminated for subsequent analysis. Peak calling for H3K27ac was performed using software MACS2 (v2.1.1) [68] in paired mode and comparing ChIP samples to input, using the false discovery rate of q < 10^-3^. H3K27me3 peaks were called by a sliding window approach to find regions enriched with H3K27me3 compared to input reads. Spike-in was used in H3K27me3 ChIP-seq for normalization because of the global loss of H3K27me3 after PRC2 loss. We discarded peaks mapped to blacklisted genomic regions identified by the ENCODE [69] [70]. Primary antibodies for ChIP are as below: anti-H3K27me3 (CST, #9733) and anti-H3K27ac (Abcam #ab4729).

The H3K27ac peaks were separated into promoters, distal and intergenic peaks as described previously [71]. They were also used for super enhancer (SE) analysis using the ROSE R package (option: -t 2500)[72] [73]. The peaks from controls and sgSUZ12 knockout samples were merged to generate a non-overlapping list of union peaks. ChIP-seq reads located to the merged peaks were calculated and used by the software DESeq2 to identify peaks with differentially modifications at the adjusted p-value < 0.05 and fold change > 2. The significantly increased or decreased peaks in the sgSUZ12 samples at promoters and non-promoter regions were subject to independent transcription factor binding motif analysis with the HOMER software (v4.7, default parameter)[74], using all peaks as background. The fold changes in these peaks were also compared to gene expression changes in the RNA-seq analysis for each of the promoter and distal peaks assigned to genes (note that one gene could have multiple peaks).

### Statistics

All statistical analyses were performed using GraphPad Prism 7. Unless otherwise noted in the figure legend, all data were shown as mean ± SEM combined with a two-tailed unpaired t-test for statistical comparisons between two groups, one-way ANOVA for more than two groups, and Log-rank (Mantel-Cox) test for survival study. Significance was defined as below: \**P*<0.05, \*\**P*<0.01, \*\*\**P*<0.001, \*\*\*\**P*<0.0001. All experiments shown were repeated in at least two independent experiments.

## Supporting information

Supplementary_Table_1_Yan_Chen_et_al

Supplementary_Table_2_Yan_Chen_et_al

Supplementary_Table_3_Yan_Chen_et_al

Supplementary_Table_4_Yan_Chen_et_al

Supplementary_Table_5_Yan_Chen_et_al

## Data availability

RNA-seq data of human MPNST tumor tissues were deposited in dbGaP with study accession: phs000792.v1.p1. Other sequencing data were deposited in the Gene Expression Omnibus: RNA-seq data of AT3 tumors in GSE179703; ATAC-seq data of M3 cells in GSE179699; ChIP-seq data of M3 cells in GSM5420909, GSM5420912, GSM5420919 and GSM5420922. All other data supporting the findings of this study are available from the corresponding authors upon reasonable request.

## Author contributions

Project conception and experimental design: J.Y., Y-D.C., P.C. and Y.C. Clinical sample collection: J.Y., Y-D.C., S.W., C.J.L., J.S., C.R.A., S.S., P.C. and Y.C. Conduction of experiments: J.Y., and Y.-D.C. performed most *in vitro* and in *vivo* experiments with assistance from A.J.P., B.G.N., E.W.P.W., M.M.R., and technical support from S.W., C.J.L., E.G., and J.S. for *in vivo* experiments. Heat-iMVA-based viral therapy: J.Y., L.D., N.Y. and Y.W. Pathology and immunohistochemistry review of clinical and murine tumor samples: C.R.A., P.C. Additional conceptual advice and experimental design and technical support for immunology experiments: M.L. L.D. and B.G.N. Computational analysis (RNA-seq, ChIP-seq, ATAC-seq, and integrative analysis): D.Y.Z., R.P.K., J.L.M., Y.L., P.M.G.J., F.Y.T., E.K., Y.C., J.Y. and P.C. Manuscript writing: J.Y., P.C. and Y.C.. All authors reviewed and edited the manuscript.

## Acknowledgments

We thank all members of the Chi and Chen laboratory for general support. We thank Drs. William Gerald and Leon (Xiaoliang) Xu for the patient derived MPNST M3 cell line. Next generation sequencing, ChIP-seq, RNA-seq and ATAC-seq were performed at the Integrative Genomics Operation (IGO)/Center for Molecular Oncology and Center for Epigenetic Research, immunohistochemistry of FFPE and MPNST samples were procured and performed at the Pathology core, and animal studies were performed at the Center of Comparative Medicine & Pathology core facility at Memorial Sloan Kettering Cancer Center (MSKCC). This work was supported by grants from the National Institute of Health (NIH) and National Cancer Institute (NCI) grants (R01 CA228216, DP2 CA174499), Department of Defense (DOD) grant (W81XWH-15-1-0124), Francis Collins Scholar NTAP, Cycle for Survival and Linn Family Discovery Fund to P.C.; the NIH/NCI grant (P50 CA217694) to P.C., S.S., C.R.A.; the NIH/NCI grants (R01 CA208100) to Y.C.; Geoffrey Beene Cancer Research Fund to P.C., M.L. and L.D., Translational Oncology Research in Oncology Training Program T32 grant (5T32CA160001-09) to A.J.P.; and the NIH grant P30 CA 008748 to Memorial Sloan Kettering Cancer Center (Core Grant).

## Declaration of interests

Memorial Sloan Kettering Cancer Center filed a patent application for the use of inactivated MVA as monotherapy or in combination with immune checkpoint blockade for solid tumors. L.D. is an author on the patent, which has been licensed to IMVAQ Therapeutics.

## References

1. Margueron, R. and D. Reinberg, The Polycomb complex PRC2 and its mark in life. Nature, 2011. 469(7330): p. 343-9.

2. De Raedt, T., et al., PRC2 loss amplifies Ras-driven transcription and confers sensitivity to BRD4-based therapies. Nature, 2014. 514(7521): p. 247-51.

3. Lee, W., et al., PRC2 is recurrently inactivated through EED or SUZ12 loss in malignant peripheral nerve sheath tumors. Nat Genet, 2014. 46(11): p. 1227–32.

4. Ernst, T., et al., Inactivating mutations of the histone methyltransferase gene EZH2 in myeloid disorders. Nat Genet, 2010. 42(8): p. 722–6.

5. Nikoloski, G., et al., Somatic mutations of the histone methyltransferase gene EZH2 in myelodysplastic syndromes. Nat Genet, 2010. 42(8): p. 665–7.

6. Ntziachristos, P., et al., Genetic inactivation of the polycomb repressive complex 2 in T cell acute lymphoblastic leukemia. Nat Med, 2012. 18(2): p. 298–301.

7. Zhang, J., et al., The genetic basis of early T-cell precursor acute lymphoblastic leukaemia. Nature, 2012. 481(7380): p. 157-63.

8. Harutyunyan, A.S., et al., H3K27M induces defective chromatin spread of PRC2-mediated repressive H3K27me2/me3 and is essential for glioma tumorigenesis. Nat Commun, 2019. 10(1): p. 1262.

9. Lewis, P.W., et al., Inhibition of PRC2 activity by a gain-of-function H3 mutation found in pediatric glioblastoma. Science, 2013. 340(6134): p. 857-61.

10. Lulla, R.R., A.M. Saratsis, and R. Hashizume, Mutations in chromatin machinery and pediatric high-grade glioma. Sci Adv, 2016. 2(3): p. e1501354.

11. Stafford, J.M., et al., Multiple modes of PRC2 inhibition elicit global chromatin alterations in H3K27M pediatric glioma. Sci Adv, 2018. 4(10): p. eaau5935.

12. Wassef, M., et al., Impaired PRC2 activity promotes transcriptional instability and favors breast tumorigenesis. Genes Dev, 2015. 29(24): p. 2547–62.

13. Prieto-Granada, C.N., et al., Loss of H3K27me3 Expression Is a Highly Sensitive Marker for Sporadic and Radiation-induced MPNST. Am J Surg Pathol, 2016. 40(4): p. 479–89.

14. Schaefer, I.M., C.D. Fletcher, and J.L. Hornick, Loss of H3K27 trimethylation distinguishes malignant peripheral nerve sheath tumors from histologic mimics. Mod Pathol, 2016. 29(1): p. 4–13.

15. Zhang, M., et al., Somatic mutations of SUZ12 in malignant peripheral nerve sheath tumors. Nat Genet, 2014. 46(11): p. 1170–2.

16. Chen, G., et al., Ezh2 Regulates Activation-Induced CD8(+) T Cell Cycle Progression via Repressing Cdkn2a and Cdkn1c Expression. Front Immunol, 2018. 9: p. 549.

17. Gray, S.M., et al., Polycomb Repressive Complex 2-Mediated Chromatin Repression Guides Effector CD8(+) T Cell Terminal Differentiation and Loss of Multipotency. Immunity, 2017. 46(4): p. 596–608.

18. He, S., et al., Ezh2 phosphorylation state determines its capacity to maintain CD8(+) T memory precursors for antitumor immunity. Nat Commun, 2017. 8(1): p. 2125.

19. Kakaradov, B., et al., Early transcriptional and epigenetic regulation of CD8(+) T cell differentiation revealed by single-cell RNA sequencing. Nat Immunol, 2017. 18(4): p. 422–432.

20. Russ, B.E., et al., Distinct epigenetic signatures delineate transcriptional programs during virus-specific CD8(+) T cell differentiation. Immunity, 2014. 41(5): p. 853–65.

21. Zhao, E., et al., Cancer mediates effector T cell dysfunction by targeting microRNAs and EZH2 via glycolysis restriction. Nat Immunol, 2016. 17(1): p. 95–103.

22. Zhang, Y., et al., The polycomb repressive complex 2 governs life and death of peripheral T cells. Blood, 2014. 124(5): p. 737–49.

23. Yang, X.P., et al., EZH2 is crucial for both differentiation of regulatory T cells and T effector cell expansion. Sci Rep, 2015. 5: p. 10643.

24. Tumes, D.J., et al., The polycomb protein Ezh2 regulates differentiation and plasticity of CD4(+) T helper type 1 and type 2 cells. Immunity, 2013. 39(5): p. 819–32.

25. De Simone, M., et al., Transcriptional Landscape of Human Tissue Lymphocytes Unveils Uniqueness of Tumor-Infiltrating T Regulatory Cells. Immunity, 2016. 45(5): p. 1135–1147.

26. DuPage, M., et al., The chromatin-modifying enzyme Ezh2 is critical for the maintenance of regulatory T cell identity after activation. Immunity, 2015. 42(2): p. 227–238.

27. Wang, D., et al., Targeting EZH2 Reprograms Intratumoral Regulatory T Cells to Enhance Cancer Immunity. Cell Rep, 2018. 23(11): p. 3262–3274.

28. Peng, D., et al., Epigenetic silencing of TH1-type chemokines shapes tumour immunity and immunotherapy. Nature, 2015. 527(7577): p. 249-53.

29. Burr, M.L., et al., An Evolutionarily Conserved Function of Polycomb Silences the MHC Class I Antigen Presentation Pathway and Enables Immune Evasion in Cancer. Cancer Cell, 2019. 36(4): p. 385–401 e8.

30. Won, H.H., et al., Detecting somatic genetic alterations in tumor specimens by exon capture and massively parallel sequencing. J Vis Exp, 2013(80): p. e50710.

31. Huang, X., et al., Targeting Epigenetic Crosstalk as a Therapeutic Strategy for EZH2-Aberrant Solid Tumors. Cell, 2018. 175(1): p. 186–199 e19.

32. Kim, J., et al., A Myc network accounts for similarities between embryonic stem and cancer cell transcription programs. Cell, 2010. 143(2): p. 313–24.

33. Mikkelsen, T.S., et al., Genome-wide maps of chromatin state in pluripotent and lineage-committed cells. Nature, 2007. 448(7153): p. 553-60.

34. Chen, D.S. and I. Mellman, Oncology meets immunology: the cancer-immunity cycle. Immunity, 2013. 39(1): p. 1–10.

35. Dunn, G.P., C.M. Koebel, and R.D. Schreiber, Interferons, immunity and cancer immunoediting. Nat Rev Immunol, 2006. 6(11): p. 836–48.

36. Parker, B.S., J. Rautela, and P.J. Hertzog, Antitumour actions of interferons: implications for cancer therapy. Nat Rev Cancer, 2016. 16(3): p. 131–44.

37. Heintzman, N.D., et al., Distinct and predictive chromatin signatures of transcriptional promoters and enhancers in the human genome. Nat Genet, 2007. 39(3): p. 311–8.

38. Roadmap Epigenomics, C., et al., Integrative analysis of 111 reference human epigenomes. Nature, 2015. 518(7539): p. 317-30.

39. Chau, V., et al., Preclinical therapeutic efficacy of a novel pharmacologic inducer of apoptosis in malignant peripheral nerve sheath tumors. Cancer Res, 2014. 74(2): p. 586–97.

40. Le, L.Q., et al., Cell of origin and microenvironment contribution for NF1-associated dermal neurofibromas. Cell Stem Cell, 2009. 4(5): p. 453–63.

41. Spranger, S. and T.F. Gajewski, Impact of oncogenic pathways on evasion of antitumour immune responses. Nat Rev Cancer, 2018. 18(3): p. 139–147.

42. Sanmamed, M.F. and L. Chen, A Paradigm Shift in Cancer Immunotherapy: From Enhancement to Normalization. Cell, 2018. 175(2): p. 313–326.

43. Dai, P., et al., Modified vaccinia virus Ankara triggers type I IFN production in murine conventional dendritic cells via a cGAS/STING-mediated cytosolic DNA-sensing pathway. PLoS Pathog, 2014. 10(4): p. e1003989.

44. Dai, P., et al., Intratumoral delivery of inactivated modified vaccinia virus Ankara (iMVA) induces systemic antitumor immunity via STING and Batf3-dependent dendritic cells. Sci Immunol, 2017. 2(11).

45. Casey, S.C., et al., MYC regulates the antitumor immune response through CD47 and PD-L1. Science, 2016. 352(6282): p. 227-31.

46. Iannello, A., et al., p53-dependent chemokine production by senescent tumor cells supports NKG2D-dependent tumor elimination by natural killer cells. J Exp Med, 2013. 210(10): p. 2057–69.

47. Koyama, S., et al., STK11/LKB1 Deficiency Promotes Neutrophil Recruitment and Proinflammatory Cytokine Production to Suppress T-cell Activity in the Lung Tumor Microenvironment. Cancer Res, 2016. 76(5): p. 999–1008.

48. Peng, W., et al., Loss of PTEN Promotes Resistance to T Cell-Mediated Immunotherapy. Cancer Discov, 2016. 6(2): p. 202–16.

49. Rakhra, K., et al., CD4(+) T cells contribute to the remodeling of the microenvironment required for sustained tumor regression upon oncogene inactivation. Cancer Cell, 2010. 18(5): p. 485–98.

50. Sai, J., et al., PI3K Inhibition Reduces Mammary Tumor Growth and Facilitates Antitumor Immunity and Anti-PD1 Responses. Clin Cancer Res, 2017. 23(13): p. 3371–3384.

51. Seiwert, T.Y., et al., Integrative and comparative genomic analysis of HPV-positive and HPV-negative head and neck squamous cell carcinomas. Clin Cancer Res, 2015. 21(3): p. 632–41.

52. Spranger, S., R. Bao, and T.F. Gajewski, Melanoma-intrinsic beta-catenin signalling prevents anti-tumour immunity. Nature, 2015. 523(7559): p. 231-5.

53. Stine, Z.E., et al., MYC, Metabolism, and Cancer. Cancer Discov, 2015. 5(10): p. 1024–39.

54. Spranger, S., et al., Tumor-Residing Batf3 Dendritic Cells Are Required for Effector T Cell Trafficking and Adoptive T Cell Therapy. Cancer Cell, 2017. 31(5): p. 711–723 e4.

55. Miao, D., et al., Genomic correlates of response to immune checkpoint therapies in clear cell renal cell carcinoma. Science, 2018. 359(6377): p. 801-806.

56. Pan, D., et al., A major chromatin regulator determines resistance of tumor cells to T cell-mediated killing. Science, 2018. 359(6377): p. 770-775.

57. Bommareddy, P.K., M. Shettigar, and H.L. Kaufman, Integrating oncolytic viruses in combination cancer immunotherapy. Nat Rev Immunol, 2018. 18(8): p. 498–513.

58. Russell, S.J. and G.N. Barber, Oncolytic Viruses as Antigen-Agnostic Cancer Vaccines. Cancer Cell, 2018. 33(4): p. 599–605.

59. Pittman, P.R., et al., Phase 3 Efficacy Trial of Modified Vaccinia Ankara as a Vaccine against Smallpox. N Engl J Med, 2019. 381(20): p. 1897–1908.

60. Volz, A. and G. Sutter, Modified Vaccinia Virus Ankara: History, Value in Basic Research, and Current Perspectives for Vaccine Development. Adv Virus Res, 2017. 97: p. 187–243.

61. Dobin, A., et al., STAR: ultrafast universal RNA-seq aligner. Bioinformatics, 2013. 29(1): p. 15–21.

62. Trapnell, C., et al., Transcript assembly and quantification by RNA-Seq reveals unannotated transcripts and isoform switching during cell differentiation. Nat Biotechnol, 2010. 28(5): p. 511–5.

63. Subramanian, A., et al., Gene set enrichment analysis: a knowledge-based approach for interpreting genome-wide expression profiles. Proc Natl Acad Sci U S A, 2005. 102(43): p. 15545–50.

64. Buenrostro, J.D., et al., Transposition of native chromatin for fast and sensitive epigenomic profiling of open chromatin, DNA-binding proteins and nucleosome position. Nat Methods, 2013. 10(12): p. 1213–8.

65. Buenrostro, J.D., et al., ATAC-seq: A Method for Assaying Chromatin Accessibility Genome-Wide. Curr Protoc Mol Biol, 2015. **109**: p. 21 29 1-21 29 9.

66. Chi, P., et al., ETV1 is a lineage survival factor that cooperates with KIT in gastrointestinal stromal tumours. Nature, 2010. 467(7317): p. 849-53.

67. Langmead, B., et al., Ultrafast and memory-efficient alignment of short DNA sequences to the human genome. Genome Biol, 2009. 10(3): p. R25.

68. Zhang, Y., et al., Model-based analysis of ChIP-Seq (MACS). Genome Biol, 2008. 9(9): p. R137.

69. Consortium, E.P., An integrated encyclopedia of DNA elements in the human genome. Nature, 2012. 489(7414): p. 57-74.

70. Carroll, T.S., et al., Impact of artifact removal on ChIP quality metrics in ChIP-seq and ChIP-exo data. Front Genet, 2014. 5: p. 75.

71. Banaszynski, L.A., et al., Hira-dependent histone H3.3 deposition facilitates PRC2 recruitment at developmental loci in ES cells. Cell, 2013. 155(1): p. 107–20.

72. Whyte, W.A., et al., Master transcription factors and mediator establish super-enhancers at key cell identity genes. Cell, 2013. 153(2): p. 307–19.

73. Loven, J., et al., Selective inhibition of tumor oncogenes by disruption of super-enhancers. Cell, 2013. 153(2): p. 320–34.

74. Heinz, S., et al., Simple combinations of lineage-determining transcription factors prime cis-regulatory elements required for macrophage and B cell identities. Mol Cell, 2010. 38(4): p. 576–89.

