## supplementary_Figures_1_to_8 for "Tumor-intrinsic PRC2 inactivation drives a context-dependent immune-desert tumor microenvironment and confers resistance to immunotherapy"

### Supplementary figures and figure legends

Supplementary Figure S1

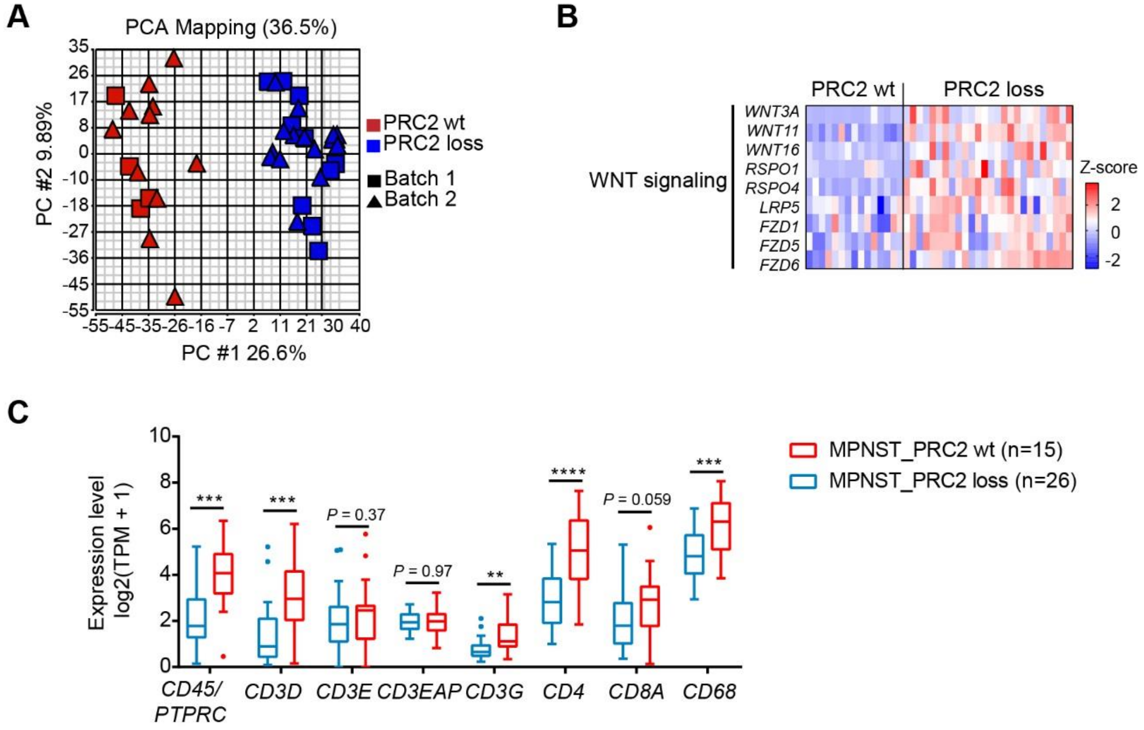


**Supplementary Figure S1. Differential gene expression between PRC2-wt and PRC2-loss human MPNSTs, Related to Figure 1. A**, Principal component analysis (PCA) of all (1st and 2nd batch) tumor transcriptomes by RNA-seq, demonstrating that the PRC2 status (PRC2-wt vs. PRC2-loss) is readily separated by principal component 1 (PC#1). **B**, Heatmap of WNT signaling pathway genes in human MPNSTs by RNA-seq. **C**, Tukey's box and whiskers plots of mRNA expression levels of immune maker genes in human MPNST by RNA-seq. TPM: transcripts per million. ****P*<0.001, *****P*<0.0001 by unpaired two-tailed t test.

Supplementary Figure S2

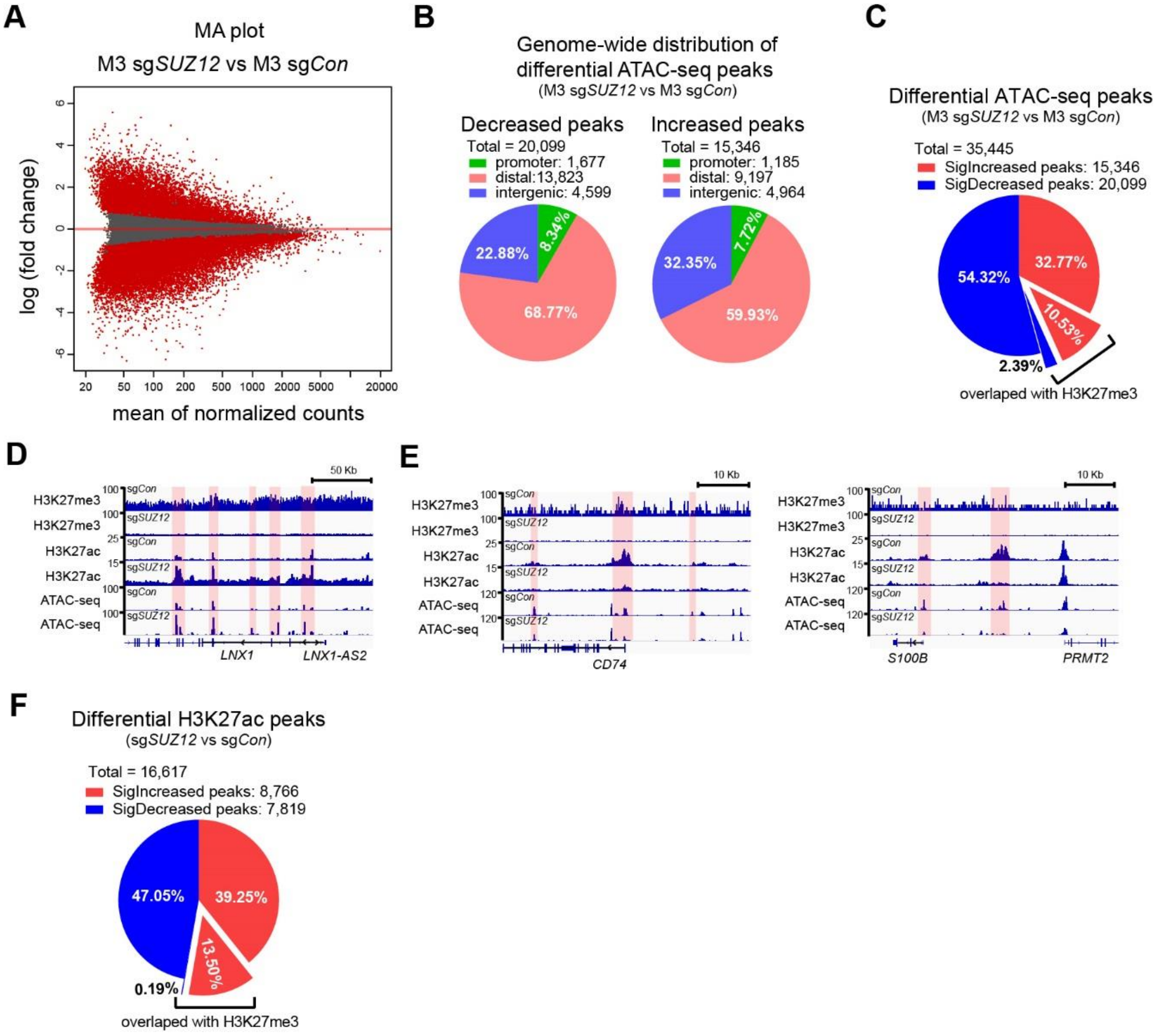


**Supplementary Figure S2. PRC2 loss reshapes the chromatin landscape, Related to Figure 3. A**, MA plot of chromatin accessibility changes by ATAC-seq comparing PRC2-loss (sg*SUZ12*) with PRC2-wt (sg*Con*) isogenic M3 cells. Red dots represent significantly changed ATAC peak (FDR $q < 0.1$, fold-change $\geq 1.5$). **B**, Genome-wide distribution of significantly decreased (left) and increased (right) chromatin accessibility sites by ATAC-seq comparing PRC2-loss (sg*SUZ12*) with PRC2-wt (sg*Con*) isogenic M3 cells (FDR $q < 0.1$, fold-change $\geq 1.5$); promoter (TSS ± 2 kb), distal regulatory (-50 kb from TSS to transcriptional end site [TES] + 5 kb) and intergenic (non-promoter, non-distal regulatory) regions. **C**, Pie chart of the overlap of the significant ATAC-seq peaks with H3K27me3 enrichment, comparing PRC2-loss (sg*SUZ12*) and PRC2-wt (sg*Con*) isogenic M3 cells. **D-E**, ChIP-seq and ATAC-seq profiles at the loci of selective genes with increased (D) and decreased (E) H3K27ac enrichment and chromatin accessibility comparing PRC2-loss (sg*SUZ12*) and PRC2-wt (sg*Con*) M3 cells. Pink highlights regions with H3K27ac enrichment changes, while yellow highlights regions with high H3K27me3 in sg*Con*. **F**, Pie chart of the overlap of the sigificant H3K27ac peaks with H3K27me3 enrichment, comparing PRC2-loss (sg*SUZ12*) and PRC2-wt (sg*Con*) isogenic M3 cells.

Supplementary Figure S3

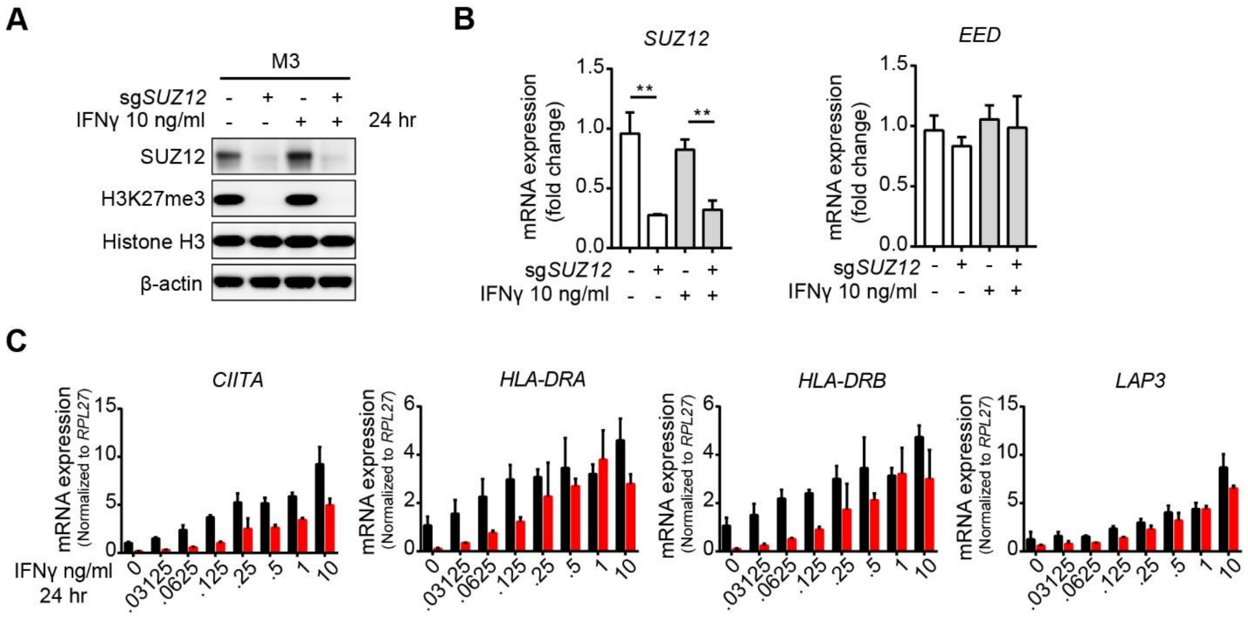


**Supplementary Figure S3. IFNγ response is diminished by PRC2 loss in M3 cells, Related to Figure 4. A**, Immunoblots of indicated protein and histone modifications in PRC2-loss (sg*SUZ12*) and PRC2-wt (sg*Con*) isogenic M3 cells stimulated with IFNγ (10 ng/ml) for 24 hours. **B**, mRNA expression levels of S*UZ12* and *EED* by qRT-PCR in PRC2-loss (sg*SUZ12*) and PRC2-wt (sg*Con*) isogenic M3 cells stimulated with IFNγ (10 ng/ml) for 24 hours. n=4, 2 in technical replicates from 2 biological replicates. **C**, IFNγ dose-dependent mRNA expression changes by qRT-PCR of IFNγ-responsive genes. n=2 in technical replicates. All error bars: mean ± SEM.

Supplementary Figure S4

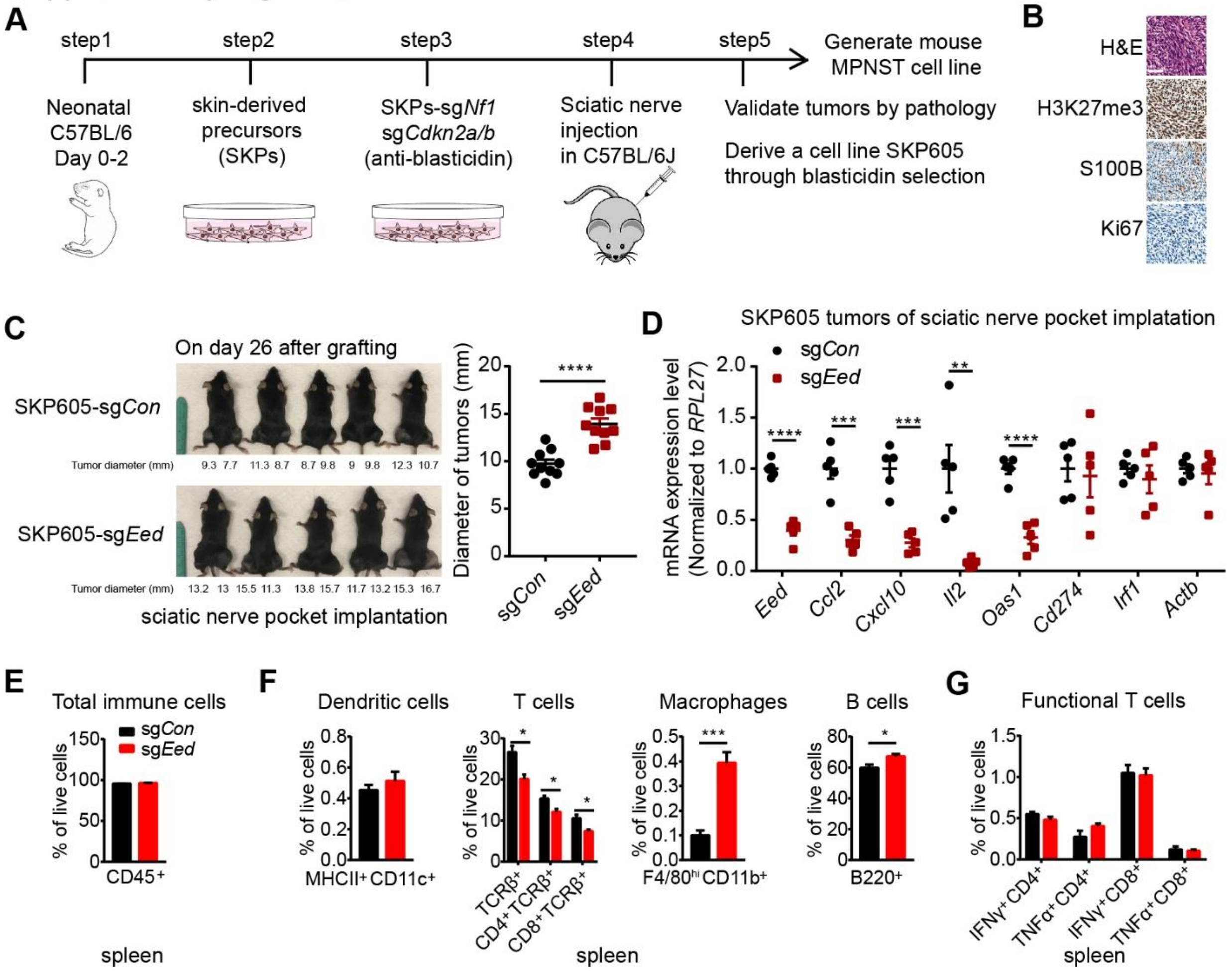


**Supplementary Figure S4. PRC2 loss promotes tumor growth and immune evasion in engineered murine MPNST model amenable for orthotopic and syngeneic transplant, Related to Figure 5. A**, A schematic of the development of a murine SKP605 MPNST model amenable for orthotopic (sciatic nerve) and syngeneic transplant in C57BL/6J mice. **B**, Representative histology and IHC of indicated proteins in orthotopically transplanted murine MPNST tumors derived from step 5 in (A). Scale bar: 50 μm. **C**. Representative images (left) and tumor sizes (right) of orthotopically and syngeneically transplanted murine PRC2-isogenic (sg*Con* vs. sg*Eed*) SKP605 MPNSTs on day 26 post-transplant. Diameter = (length + width + height)/3. n=10 tumors for each group. ****$P$<0.0001 by unpaired two-tailed t test. **D**, Relative mRNA expression change of indicated genes by qRT-PCR in PRC2-loss (sg*Eed*) and PRC2-wt (sg*Con*) SKP605 tumors in (C). n=5 tumors for each group. **$P$<0.01, ***$P$<0.001, ****$P$<0.0001 by unpaired two-tailed t test. **E-F**, Percentage of CD45$^+$ (E) and subpopulations (F) of immune cells, including dendritic cells (B220$^-$F4/80$^{lo}$MHCII$^+$CD11c$^+$), T cells (TCRβ$^+$), macrophages (B220$^-$F4/80$^{hi}$CD11b$^+$) and B cells (B220$^+$), of total live cells in the spleen of PRC2-loss (sg*Eed*) and PRC2-wt (sg*Con*) SKP605 tumors. n=5 spleens for each group. *$P$<0.05, ***$P$<0.001 by unpaired two-tailed t test. **G**, The percentage of IFNγ$^+$CD4$^+$ or IFNγ$^+$CD8$^+$ T cells of total live cells in the spleen of PRC2-isogenic SKP605 tumors. n=5 spleens for each group. All error bars: mean ± SEM.

Supplementary Figure S5

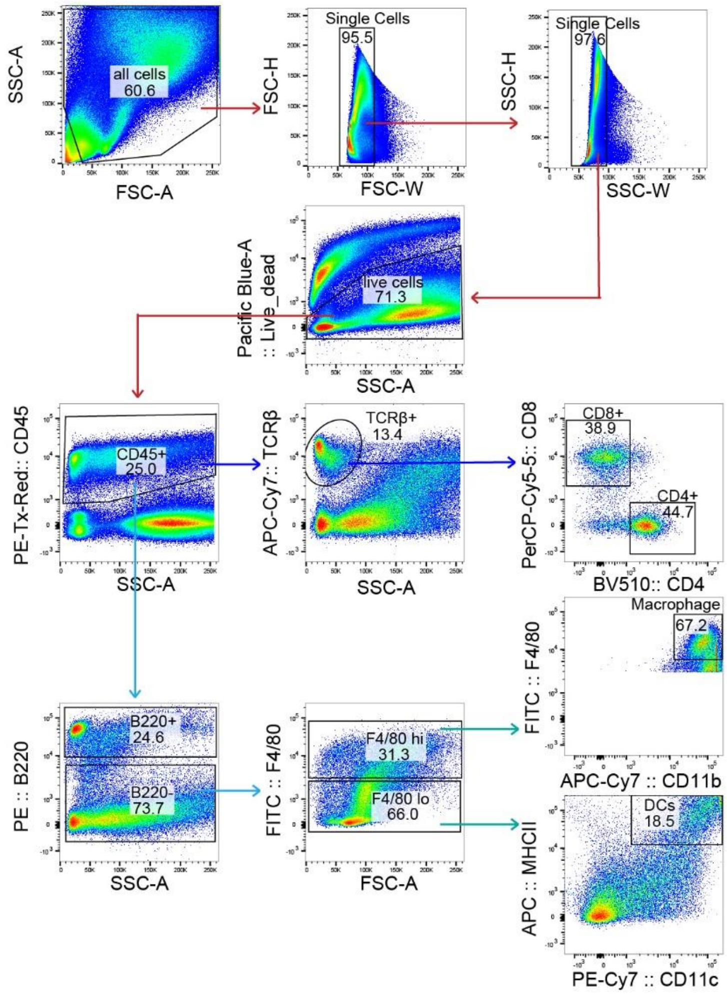


**Supplementary Figure S5. Gating strategies of multi-color flow cytometry for immune profile, Related to Figure 5.** Representative example of CD45, B220, F4/80, MHCII, CD11c, CD11b, TCRβ, CD4, CD8 staining to show gating skills (gated for total immune cells, dendritic cells, T cells, macrophage and B cells) for different immune subpopulations by FACS.

Supplementary Figure S6

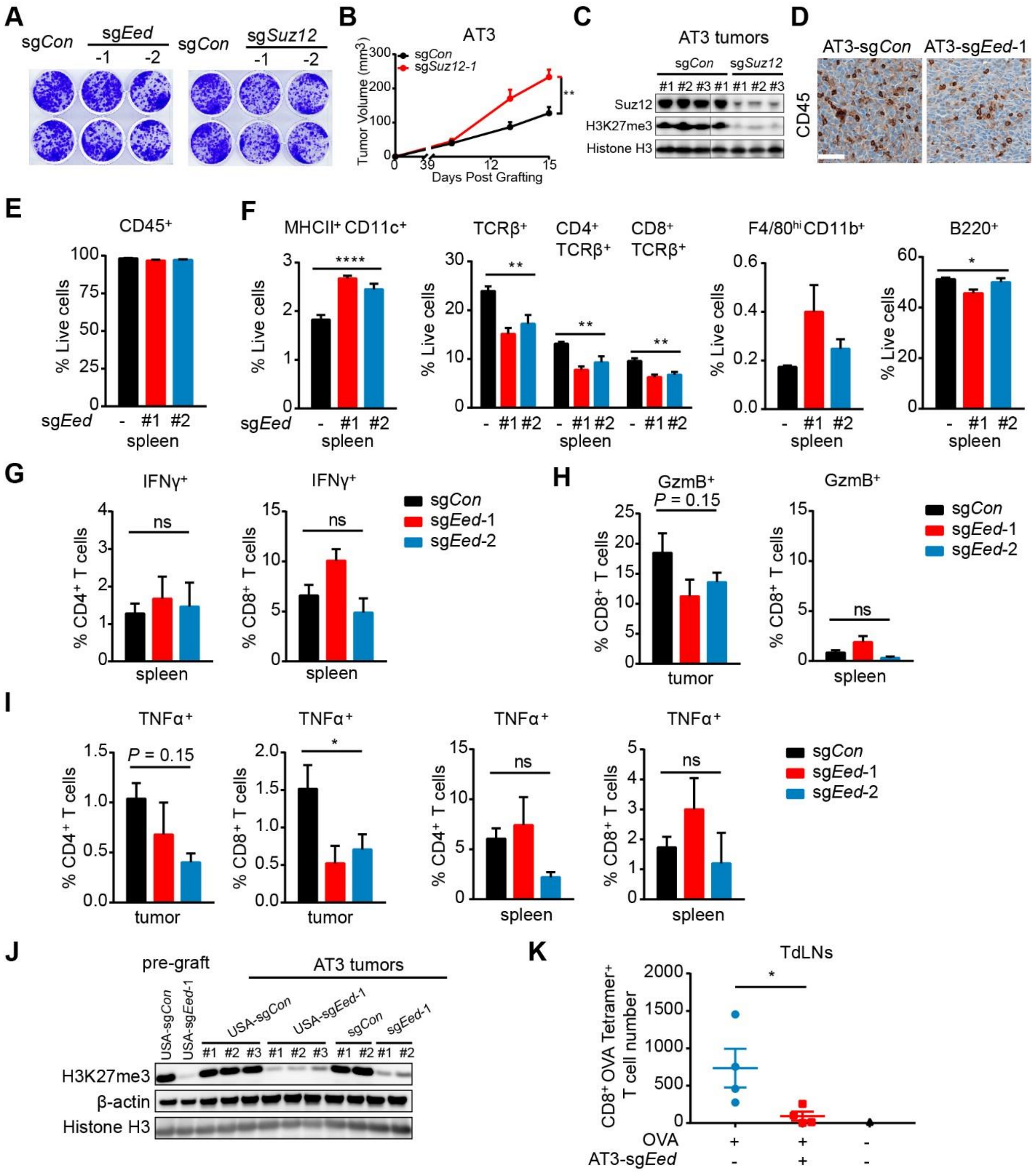

**Supplementary Figure S6. PRC2 loss promotes tumor growth and immune evasion in murine breast cancer model, Related to Figure 6. A**, Clonogenic assays of PRC2-wt (sg*Con*) and PRC2-loss (sg*Eed* or sg*Suz12*) AT3 cells. n=2 replicates for each group. **B**, Growth curves of PRC2-isogenic (sg*Con* vs. sg*Suz12*) AT3 syngeneically transplanted tumors in the mammary fat pads of C57BL/6J mice. n=10 tumors for each group. $^{**}P<0.01$ by unpaired two-tailed t test. **C**, Immunoblots of indicated proteins and histone modifications, confirming PRC2-status (PRC2-wt [sg*Con*] vs. PRC2-loss [sg*Suz12*]) in representative syngeneically transplanted AT3 tumors. **D**, Representative IHC of $CD45^+$ immune cells in PRC2-isogenic AT3 tumors. Scale bar: 50 μm. **E-F**, Percentage of $CD45^+$ immune cells (E) and subpopulations of immune cells (F), including dendritic cells ($B220^-F4/80^{lo}MHCII^+CD11c^+$), T cells ($TCR\beta^+$), macrophages ($B220^-F4/80^{hi}CD11b^+$) and B cells ($B220^+$), in the spleens of PRC2-isogenic AT3 tumors. n=5 spleens for each group. $^*P<0.05$, $^{**}P<0.01$, $^{****}P<0.0001$ by unpaired two-tailed t test. **G**, Percentage of $IFN\gamma^+$ cells in $CD4^+$ and $CD8^+$ T cells in the spleens of PRC2-isogenic AT3 tumors. n=5 spleens for each group. **H-I**, Percentage of $GzmB^+$ (H) and $TNF\alpha^+$ (I) cells in $CD4^+$ or $CD8^+$ T cells in PRC2-isogenic AT3 tumors (left) and spleens (right). n= 5-10 for each cohort. J, Representative immunoblots of indicated proteins and H3K27me3 in pre-graft AT3 cells and explanted AT3 tumor grafts. K, Absolute cell numbers of MHCI-OVA $tetramer^+$ $CD8^+$ T cells in TdLNs 14 days after syngeneic transplant of PRC2-isogenic AT3 cells in the mammary fad pads of C57BL/6J mice. TdLNs of $OVA^+$ tumors: n=4 for each group, TdLN of sgCon and $OVA^-$ tumor: n=1, $^*P<0.05$ by unpaired two-tailed t test. All error bars: mean ± SEM.

Supplementary Figure S7

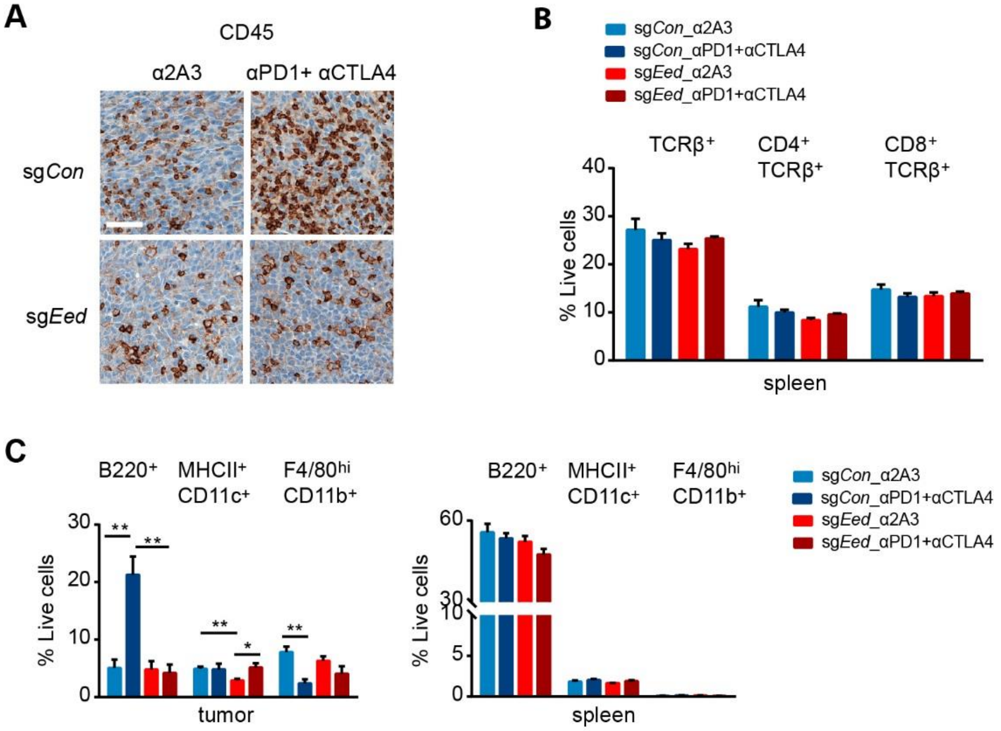


**Supplementary Figure S7. PRC2 loss confers primary resistance to immune checkpoint blockade (ICB) in murine syngeneic transplant mammary tumor models, Related to Figure 7. A**, Representative IHC of $CD45^{+}$ immune cells in PRC2-isogenic AT3 tumors treated with ICB (αPD1+αCTLA4) or control antibodies (α2A3). Scale bar: 50 µm. **B**, Percentage of T cells ($TCR\beta^{+}$), $CD4^{+}$ and $CD8^{+}$ subpopulation of T cells in the spleens of mice treated with ICB and control antibodies. n=5 for each cohort. **C**, Percentage of subpopulations of immune cells, including dendritic cells ($B220^{-}F4/80^{lo}MHCII^{+}CD11c^{+}$), T cells ($TCR\beta^{+}$), macrophages ($B220^{-}F4/80^{hi}CD11b^{+}$) and B cells ($B220^{+}$), in PRC2-isogenic AT3 tumors (left) and spleens (right) of mice treated with ICB and control antibody. n=5 for each cohort. *P<0.05, **P<0.01 by unpaired two-tailed t test. All error bars: mean±SEM.

Supplementary Figure S8

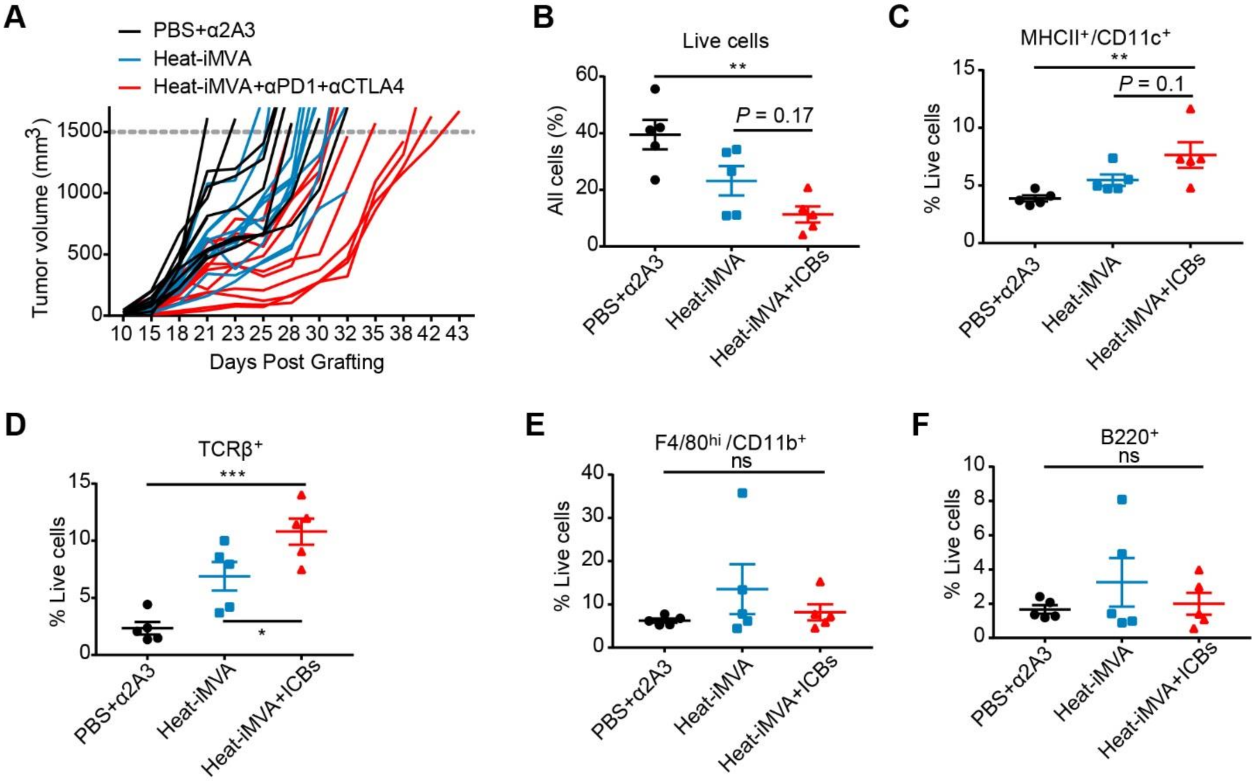


**Supplementary Figure S8. Intratumoral injection of heat-iMVA combined with anti-PD1 and anti-CTLA4 inhibits the tumor growth of PRC2-loss AT3 (sg*Eed*) tumors, Related to Figure 8. A,** Tumor growth curve of individual tumor under various treatment as indicated. Dose of treatment: $8 \times 10^7$ plaque-forming units heat-iMVA (intratumoral injection), 250 µg anti-PD1 and 200 µg anti-CTLA4, or combination per tumor. **B-F**, Percentage of live cells in all cells (B); percentage of dendritic cells ($B220^-F4/80^{lo}MHCII^+CD11c^+$) (C), T cells ($TCR\beta^+$) (D), macrophages ($B220^-F4/80^{hi}CD11b^+$) (E), and B cells ($B220^+$) (F) of total live cells in AT3 (sg*Eed*) tumors under various treatment as indicated. ICBs: anti-PD1 + anti-CTLA4. **B-F**: n=5 for each cohort. $^{**}P<0.01$, $^{***}P<0.001$ by one-way ANOVA among all three groups, and by unpaired two-tailed t test between heat-iMVA and heat-iMVA + ICB. All error bars: mean±SEM.
